# An open long non-coding RNA foundation model for generalizable prediction

**DOI:** 10.64898/2026.11.20.768117

**Authors:** Ao Chang, Heqin Zhu, Ruifeng Li, Haobin Chen, Chenxi Wang, Fenghe Tang, Qingsong Yao, Peng Xiong, S. Kevin Zhou

## Abstract

Long non-coding RNAs (lncRNAs) play essential roles in transcriptional regulation and chromatin organization, yet predicting their functional properties from sequence remains challenging because informative signals span both local motifs and long-range base pairing across kilobase-scale transcripts. Current RNA foundation models are restricted to short sequences and existing lncRNA prediction models narrowly focus on isolated tasks, leaving a dedicated foundation model notably absent. Here we present LncRFM, a dedicated lncRNA foundation model, supported by the LncCorpus pretraining data and the LncBench evaluation suite, which stand for a systematic framework to decode lncRNAs. For foundation model pretraining, we develop a hybrid bidirectional state-space encoder trained with a structure-guided masking strategy to effectively capture both local sequence patterns and long-range contexts. For pretraining data, we contribute over 9.6 million lncRNA sequences to assemble LncCorpus, a pretraining corpus of 24.7 million sequences. For benchmarking evaluation, we construct a comprehensive benchmark, LncBench, comprising 15 datasets and totaling over 1,010,000 sequences across 10 prediction tasks: six core lncRNA functional and annotation tasks and four structural and regulatory RNA assay tasks. Evaluated against seven RNA foundation models under a unified frozen-encoder protocol, our LncRFM achieves the state-of-the-art primary-metric scores on 11 of the 13 datasets that allow downstream adaptation. On two external generalization benchmarks evaluated without further adaptation, LncRFM matches the best structure-specialized baseline in F1 (0.141) and achieves the highest AUROC (0.962) on lncRNA contact prediction, while maintaining competitive cross-species coding-potential transfer (F1 of 0.374) under severe multi-kingdom domain shift. To benefit the scientific community, we make all datasets, source code, and pretrained models publicly available.

## 1 Introduction

Long non-coding RNAs (lncRNAs) participate in transcriptional regulation, chromatin organization and cellular homeostasis, making them critical biomarkers and therapeutic targets [1, 2]. Because experimental characterization of their subcellular localization, molecular interactions and disease associations remains resource-intensive, sequence-based prediction plays a vital role in prioritizing transcripts for functional study. However, decoding lncRNA functional properties from primary sequence remains challenging: unlike protein-coding transcripts, lncRNAs often exhibit weak primary-sequence conservation while preserving function through modular elements acting across multiple scales within kilobase-long transcripts. Locally, short sequence motifs govern RNA-binding protein (RBP) recognition and nuclear retention [3–6]; globally, distant nucleotides pair to form secondary structures and longrange domains that modulate site accessibility and molecular scaffolding [7, 8].

Existing computational paradigms struggle to reconcile these multiscale requirements. On the one hand, self-supervised RNA foundation models [9–15] and structure-informed pretraining frameworks [16, 17] deal primarily with short sequences and general RNA corpora, in which full-length lncRNAs are underrepresented. Because the attention architectures they rely on are constrained by context length and quadratic complexity, an lncRNA transcript averaging 1.2–3.8 kb must be truncated or split across input windows, severing the long-range pairing signals that shape its function. Consequently, models trained on general RNA corpora or short regulatory sequences show limited accuracy on lncRNA subcellular localization and protein-binding prediction even under matched evaluation conditions [18, 19], while methods that require predicted structure as input incur a separate folding step for each transcript at inference. On the other hand, existing lncRNA predictors and specialized domain models narrowly focus on isolated tasks. Specifically, task-specific lncRNA tools remain compartmentalized into standalone models for subcellular localization [20, 21], coding potential assessment [22–24], RNA–protein interaction [25, 26], or disease association prediction [27, 28]. Similarly, although incorporating structural priors or tailored architectures has advanced specialized structural and interaction modelling tasks [29–33], these supervised predictors typically rely on hand-crafted sequence features or task-specific training from scratch, failing to provide transferable representations across the diverse functional landscape of kilobase-scale lncRNAs. Thus, the remaining gap is a dedicated foundation model for prediction across lncRNA functional tasks, combining local sequence motifs with base-pairing information. Such foundation model is pretrained on a large amount of data to learn general patterns and provide reusable representations, which would support localization, protein-binding and functional annotation tasks and help prioritize lncRNAs for experimental study. Evaluating this capability requires a benchmark centred on lncRNA functional properties and transcript length, complementing existing RNA benchmarks [18, 19]. In sum, bridging the above gap requires overcoming two core challenges: establishing a massive lncRNA-enriched pretraining corpus alongside a unified multitask benchmark, and learning an encoder that captures both local patterns and kilobase-range structural contexts from sequence alone.

In this paper, we develop LncRFM, a dedicated lncRNA foundation model, supported by the LncCorpus pretraining resource and the LncBench evaluation suite. First, for model pretraining, we propose a hybrid bidirectional state-space encoder [34] trained with structure-guided pretraining mechanisms to capture both local motifs and long-range contexts. Specifically, we resolve local motifs of varying length via a multiscale convolutional stem before propagation through a hybrid backbone of ten bidirectional state-space blocks interleaved with two self-attention layers, which processes kilobase-scale transcripts with near-linear complexity beyond the context limits of pure attention-based encoders. We leverage predicted base pairing to guide two synergistic mechanisms: structure-guided joint masking draws paired nucleotides together into the reconstruction target, and pair-gated fusion weights forward and reverse contextual states by pairing status. Because structural guidance is confined to pretraining, downstream adaptation and inference operate directly on raw sequence without a folding step: for tasks supporting adaptation, the pretrained encoder remains frozen while only a lightweight multi-branch bottleneck adapter and task head are updated. Second, for data curation, we contribute over 9.6 million lncRNA transcripts extending up to 2,050 nt and assemble LncCorpus, a structure-annotated pretraining corpus of 24.7 million RNA sequences bridging short regulatory RNAs and multi-kilobase transcripts. To rigorously evaluate representation learning across this regime, we construct LncBench, a comprehensive benchmark comprising 10 prediction tasks and 15 datasets, totaling over 1,010,000 sequences. The 10 LncBench tasks encompass six core lncRNA functional and annotation tasks — (i) subcellular localization, (ii) cytoplasmic-to-nuclear relative concentration index (RCI) regression, (iii) coding potential prediction, (iv) official variation-record status annotation (Disease), (v) multi-context expression capacity prediction (Expression), and (vi) RNA–protein binding prediction — and four structural and regulatory RNA assay tasks — (vii) RNA contact prediction, (viii) dimethyl sulfate (DMS) reactivity prediction, (ix) mean ribosome load (MRL) prediction, and (x) splice-site recognition. The 15 datasets include five newly formulated task datasets curated in this study from primary public databases [35–37] and ten datasets adapted or incorporated from established resources [10, 13, 14, 38]. Among them, 13 datasets serve as internal benchmarks for downstream adaptation and 2 are held out for external generalization evaluation.

Evaluations across LncBench demonstrate that LncRFM achieves strong transferable representation learning across functional and structural regimes. Under a frozen-encoder protocol against seven representative RNA foundation models, LncRFM achieves the highest primary-metric score on 11 of the 13 adaptation datasets (Fig. 2a–m) and leads or ties for the highest unrounded score on 49 of 75 evaluated dataset–metric combinations (Fig. 3a,b), delivering consistent gains across subcellular localization, RNA–protein interaction, annotation, and structural probing tasks. Specifically, LncRFM attains the highest primary-metric scores on eight of the nine core lncRNA datasets: AUROC of 0.757, 0.791, and 0.980 on POSTAR3-UPF1, Disease, and GENCODE-Coding, respectively; Macro-AUPRC of 0.498 on Expression; F1 of 0.58 on lncATLAS-Localization; Micro-F1 of 0.76 and 0.77 on iLoc-lncRNA D1 and D2; and Pearson correlation *r* of 0.403 on lncATLAS-RCI regression (Fig. 2a–h). On the remaining core dataset, RNAErnie-lncRNA, LncRFM ranks second in AUROC behind LncRNA-BERT (Fig. 2l). Length-stratified boot-strap analyses (Fig. 3d) show that while attention-based baselines are constrained by shorter context windows, LncRFM maintains high rank-1 winning shares into the multi-kilobase regime (*>* 2, 050 nt), reaching 99.65%–100.00% on Expression, Disease, lncATLAS-RCI, GENCODE-Coding, and lncATLAS-Localization, and nearly tripling its lncATLAS-Localization F1 margin from +0.024 (1, 024 nt) to +0.068 (*>* 2, 050 nt). Notably, on fine-grained structural and translational assays, while operating directly on primary sequence without secondary-structure folding at inference, LncRFM also surpasses the structure-pretrained structRFM and sequence baselines, raising base-pair contact F1 from 0.12 to 0.16 on mRNA-Structure, squared Pearson correlation *r*^2^ from 0.28 (LncRNA-BERT) to 0.35 on 5UTR-MRL, and reactivity correlation from 0.66 to 0.67 on mRNA-DMS (Fig. 2i–k). In direct transfers of the internally adapted checkpoints to the two held-out external benchmarks without further tuning (Fig. 5), LncRFM closely matches the top structure-specialized baseline on external human lncRNA contact prediction (lncRNA-Structure, F1 of 0.141 versus 0.141 for structRFM, or 0.140621 versus 0.141048 unrounded, with a leading AUROC of 0.962), while achieving an F1 score of 0.374 (behind RNA-FM at 0.429) under pronounced cross-kingdom domain shift on multi-species coding-potential assessment (RNAChallenge). Systematic ablation analyses (Fig. 6) together with prediction diagnostics (Fig. 4) further showed that the full model outperformed the base encoder on 10 of 13 adaptation tasks and achieved the highest score among five configurations on eight tasks, including an increase in Disease AUROC from 0.765 to 0.791 and in iLoc-lncRNA D2 Micro-F1 from 0.715 to 0.770, whereas compact variants remained preferable on localization and splice-site prediction.

**Fig. 1:**
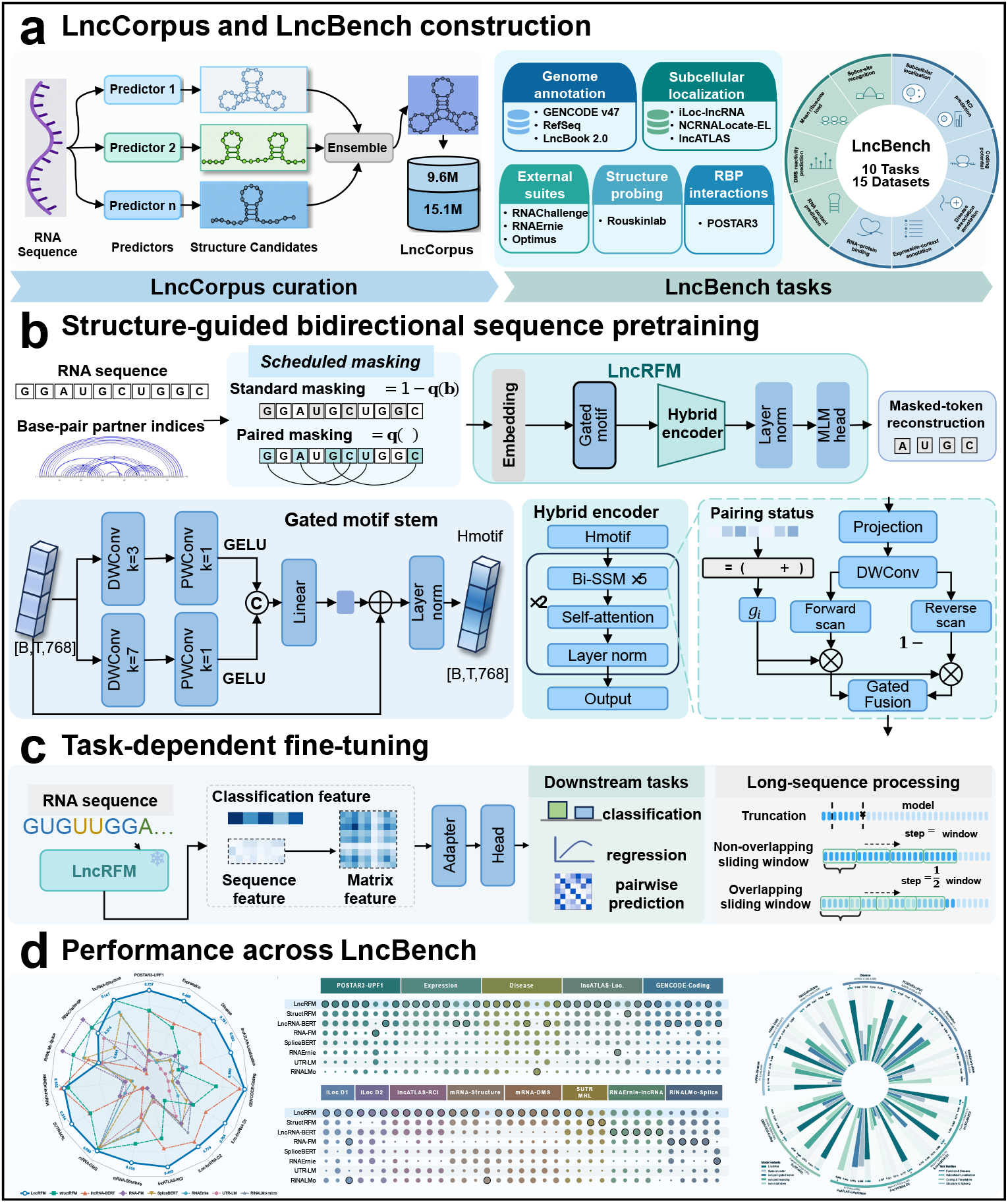
Overview of LncCorpus, LncBench and the LncRFM framework. **a**, Construction of the 24.7-million-sequence LncCorpus pretraining database (left) and the 15-dataset LncBench benchmark (right). **b**, Pairing-guided pretraining: predicted base-pair partners guide joint masking, and binary pairing status (*m_i_*) controls pair-gated fusion of forward and reverse state-space (Bi-SSM) states after multiscale motif extraction. **c**, Sequence-only adaptation: directional states are averaged without structural inputs, and frozen encoder states pass through lightweight adapters and task heads across sequence, nucleotide and pairwise tasks. **d**, Benchmark-wide evaluation overview: primary metrics across 15 datasets (left), multi-metric relative-score profiles (middle) and module ablations across 13 adapted tasks (right).

**Fig. 2:**
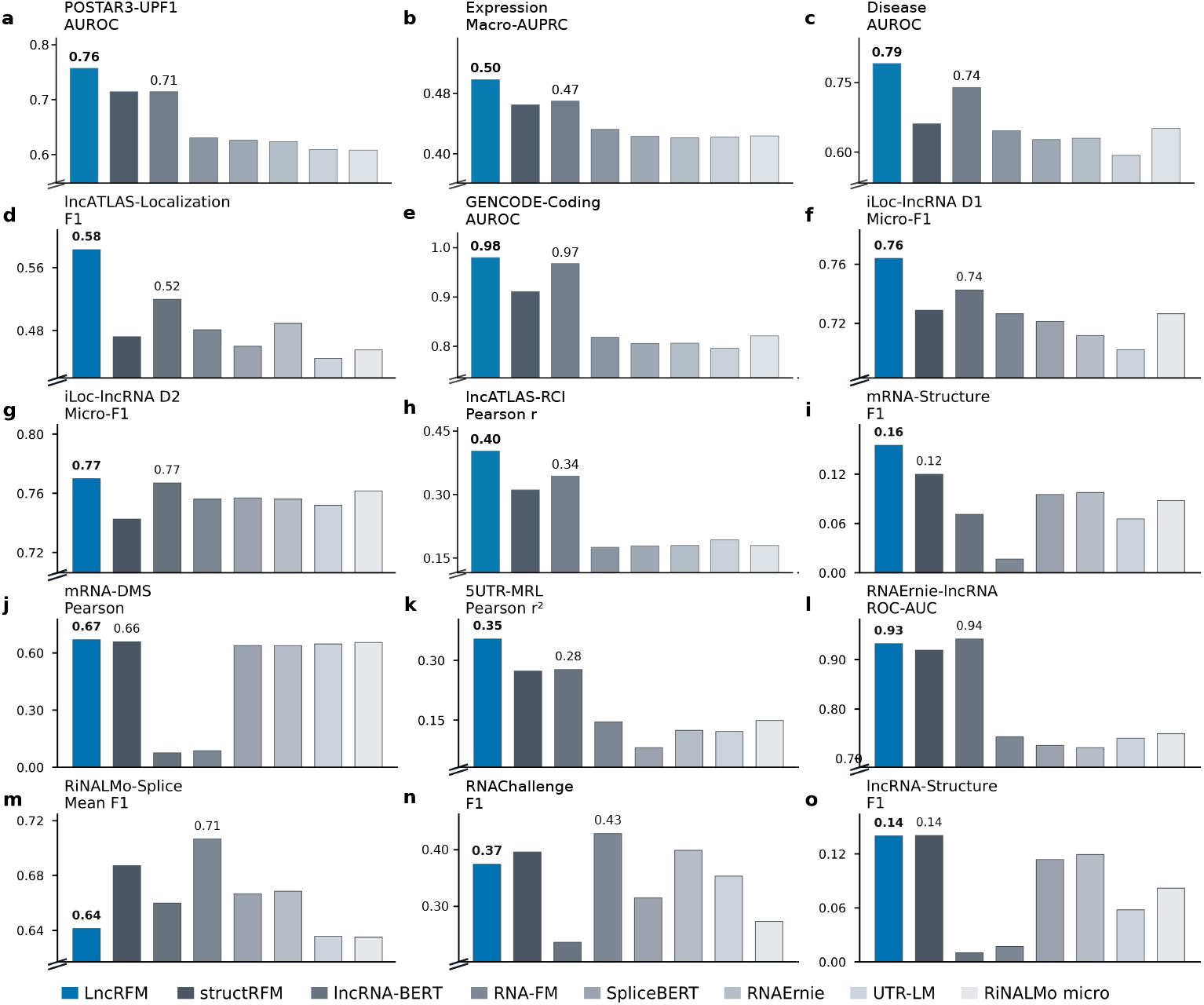
Primary-metric comparison of eight RNA encoders across 15 LncBench datasets. a–m, Thirteen adapted datasets; RiNALMo-Splice (**m**) reports mean donor/acceptor F1. **n,o**, External evaluations on RNAChallenge and lncRNA-Structure without further adaptation. Numeric labels identify LncRFM and the strongest baseline (displayed to two decimal places). Panel scales differ; diagonal marks denote non-zero lower limits. Extended Data Tables 1 and 6 give partition counts and primary-metric scores, respectively.

**Fig. 3:**
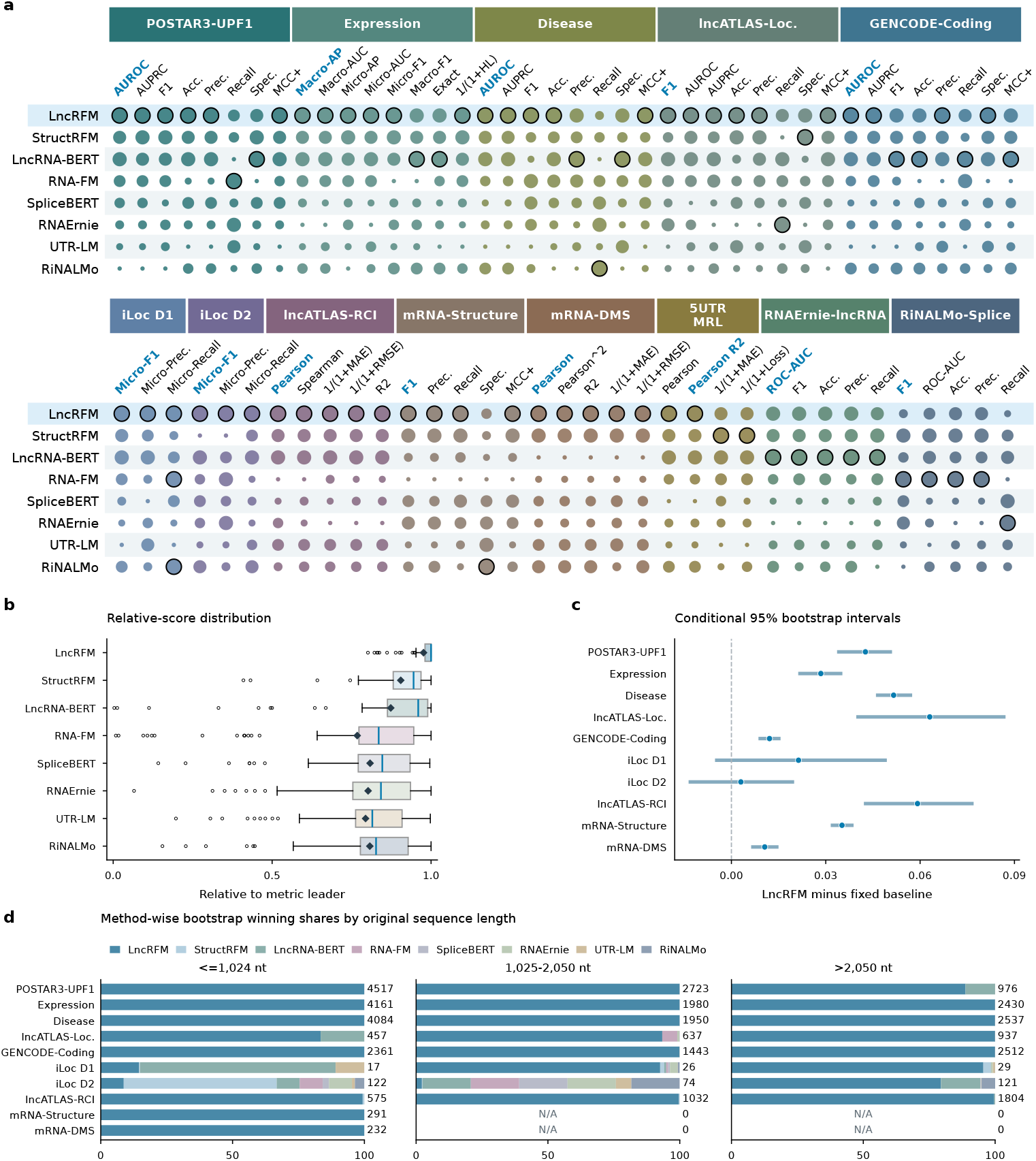
Performance across metrics, bootstrap resamples and sequence lengths. **a**, Metric profiles across 13 adapted datasets. Dot area encodes within-metric rank from unrounded scores; colour identifies datasets. Black outlines mark only unrounded maxima, including ties. Primary-metric labels are blue and bold; the LncRFM row has a light-blue background. Error metrics use 1*/*(1 + error) and MCC uses (MCC + 1)*/*2. **b**, Relative-score distributions over 75 dataset–metric combinations per model: each unrounded transformed score is divided by the corresponding metric’s unrounded maximum. Boxes, interquartile ranges (IQRs); centre lines, medians; diamonds, means; whiskers, extremes within 1.5 *×* IQR of box limits; circles, outliers. **c**, Primary-metric differences from the strongest baseline fixed across bootstrap replicates for each of ten core datasets, on the original metric scale; bars show 95% percentile intervals from 2,000 paired record bootstraps. **d**, Winning shares across 2,000 paired bootstraps within each transcript-length bin; ties split credit equally, counts appear at right and N/A denotes empty bins.

**Fig. 4:**
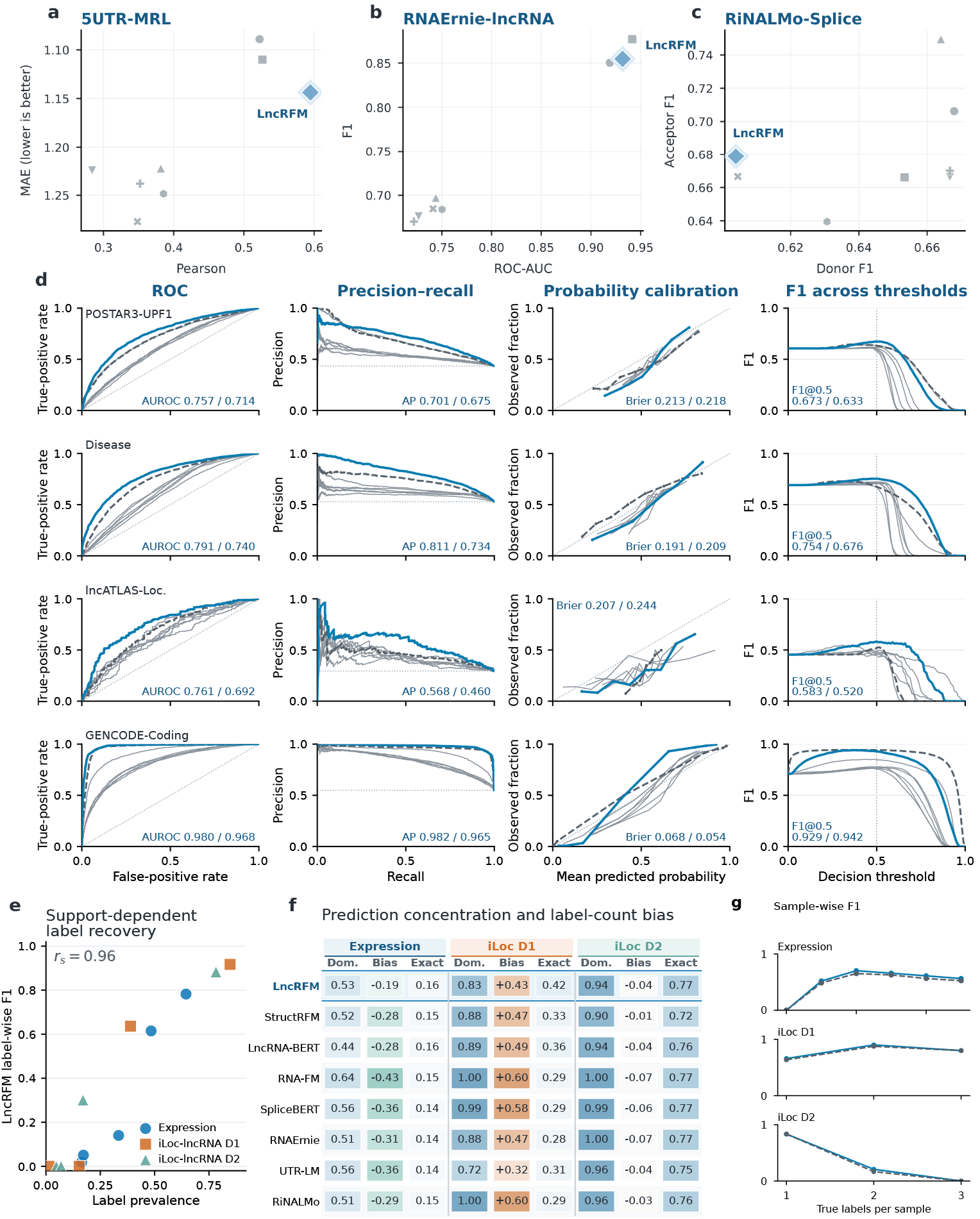
Prediction profiles across classification, regression and multilabel tasks. a–c, Joint metric profiles for 5UTR-MRL (reversed MAE axis), RNAErnie-lncRNA and RiNALMo-Splice; diamonds identify LncRFM. **d**, ROC, precision–recall, calibration and F1-threshold curves for POSTAR3-UPF1, Disease, lncATLAS-Localization and GENCODE-Coding. LncRFM is blue, the strongest primary-metric baseline is dashed grey and other methods are light grey. Numeric pairs give LncRFM/baseline AUROC, average precision, Brier score or F1 at threshold 0.5. Calibration uses eight quantile bins; reference lines show ROC/calibration equality, positive prevalence and threshold 0.5, respectively. **e**, LncRFM label-wise F1 versus prevalence across Expression and iLoc-lncRNA D1/D2 (13 labels; Spearman *r_s_* = 0.96). **f**, Dominant predicted label-set fraction (Dom.), mean predicted-minus-observed label-count bias and exactmatch fraction. LncRFM’s Dom. of 0.94 on iLoc D2 denotes predictions containing only Exosome in 94% of records. **g**, Mean sample-wise F1 by observed label count for LncRFM and the strongest baseline; Expression bands show conditional 95% record-bootstrap intervals.

**Fig. 5:**
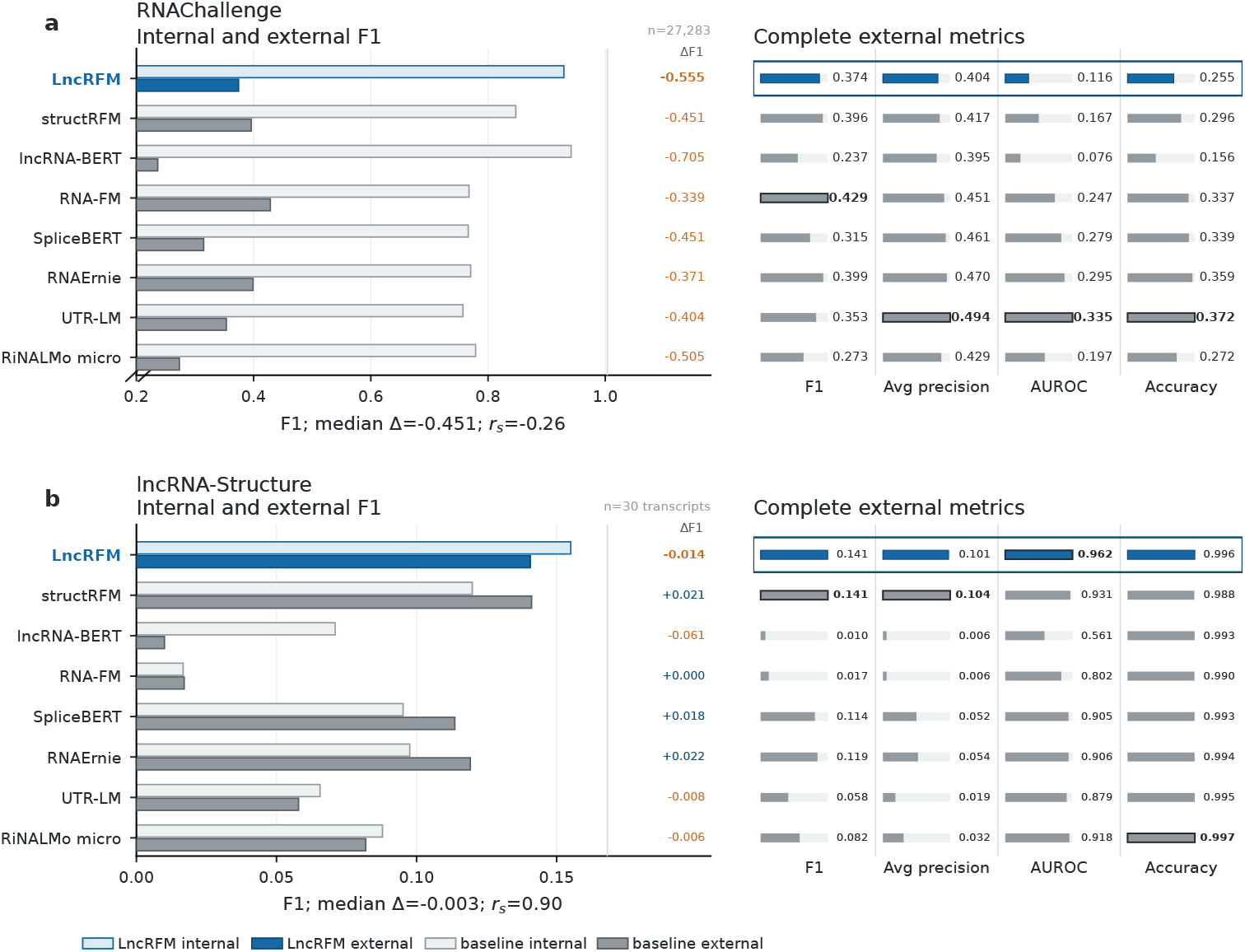
Internal and external performance of adapted checkpoints. **a,b**, Eight-model evaluations on RNAChallenge (*n* = 27, 283 records) and lncRNA-Structure (*n* = 30 transcripts), respectively. Left, internal (pale) and external (solid) F1 for LncRFM (blue) and baselines (grey); ΔF1 is external minus internal. Annotations give median ΔF1 and Spearman *r_s_* across models. The RNAChallenge F1 axis starts above zero. Right, external F1, average precision, AUROC and accu_-_racy; microbars are scaled to each metric’s leader and boldface marks the highest unrounded score. Internal reference scores correspond to the human GENCODE-Coding (**a**) and mRNA-Structure (**b**) source cohorts from which the transferred checkpoints were adapted.

**Fig. 6:**
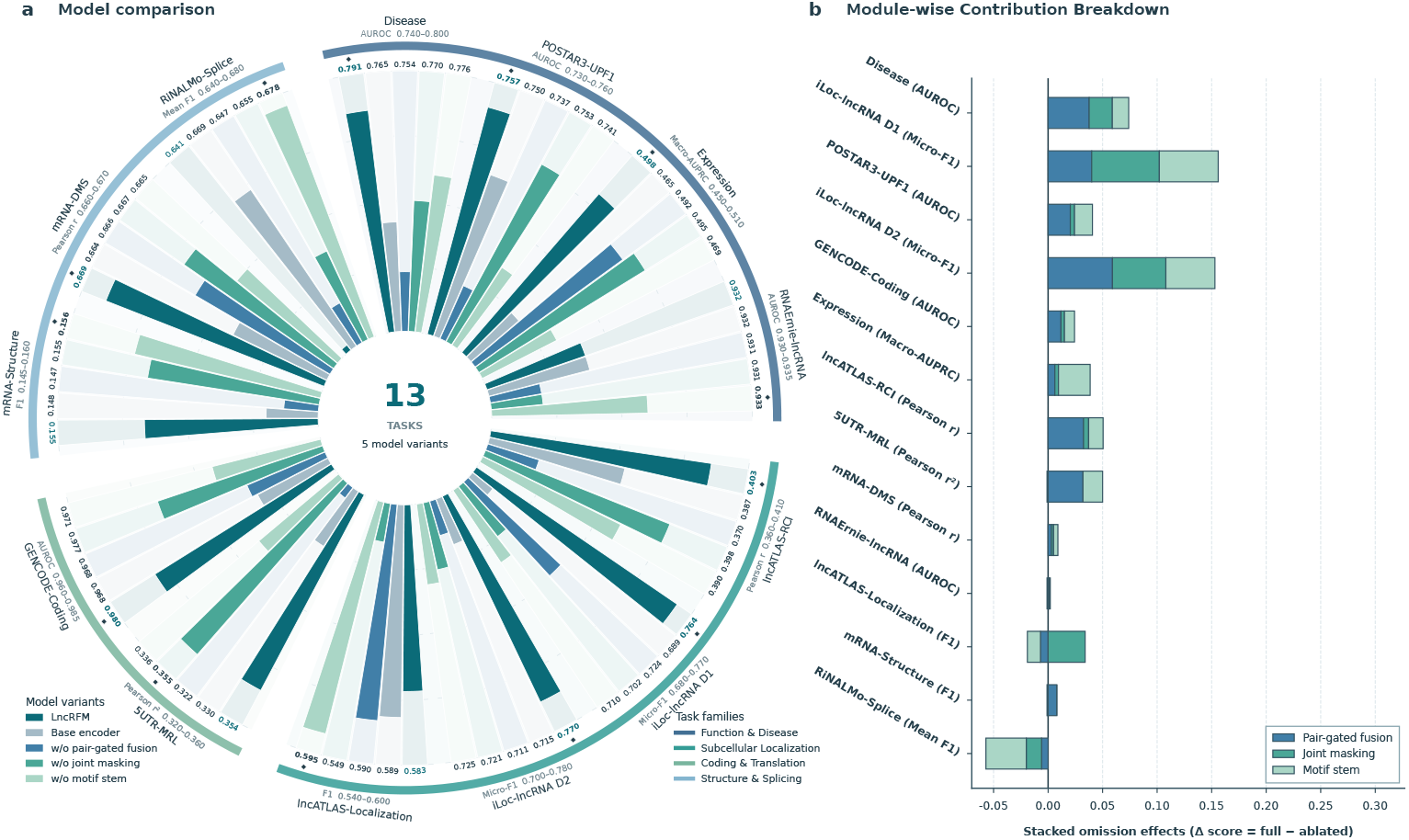
Task-dependent contributions of motif extraction and pairing-guided pretraining. The base encoder lacks the local motif stem, pairing-partner joint masking and pair-gated fusion, whereas each leave-one-out variant removes only the named component. All variants are evaluated on the identical LncBench Test cohorts used in Figs. 2–4 (Extended Data Table 1). **a**, Primary-metric scores across 13 adaptation tasks. Numeric labels are displayed to three decimal places; bold labels and diamond markers indicate the highest unrounded score for each task. **b**, Signed omission effects, Δ = *S*_full_ *− S*_ablated_, calculated from the unrounded scores. Positive values indicate a lower score after component omission, whereas negative values indicate that the ablated configuration scored higher.

To facilitate broad adoption and accelerate functional lncRNA discovery, we make the LncCorpus resource, the LncBench evaluation suite, source code, and pretrained LncRFM weights publicly available.

## 2 Results

### 2.1 The LncCorpus bridging short RNAs and kilobase-scale lncRNAs

Capturing long-range sequence–structure dependencies in non-coding transcripts requires pretraining data that span the multi-kilobase length regime. Building upon our structure-aware MUSES corpus [17]—which originally contained approximately 21.5 million short RNA sequences (512 nt) annotated via ensemble folding methods—we excluded sequences shorter than 200 nt and retained 15.1 million sequences (200–512 nt), and incorporated over 9.6 million additional lncRNA sequences extending up to 2,050 nt. Together, they are assembled into LncCorpus, a final pretraining corpus of 24.7 million sequences (Fig. 1a). Because conventional thermodynamic folding becomes computationally prohibitive across millions of kilobase-long transcripts, we annotated secondary structures for the long-sequence expansion using LinearFold [39], a linear-time folding algorithm that preserves long-range base-pairing accuracy. By integrating multi-method structural priors for short RNAs with LinearFold-derived pairing topologies for long transcripts, LncCorpus expands the maximum pretraining context fourfold (from 512 to 2,050 nt) and provides a structural foundation for modelling kilobase-scale RNA grammar.

### 2.2 The LncBench for multi-scale functional and structural evaluation

To rigorously evaluate whether pretrained representations generalize across diverse biological readouts, we constructed LncBench, a downstream benchmark comprising 15 datasets across 10 prediction tasks curated from public transcriptomic databases, experimental assays, and established benchmarks (Fig. 1a; Extended Data Table 1). Biologically, LncBench is organized into two complementary tiers. The first tier targets six core lncRNA functional and annotation properties across transcripts averaging 1.2 to 3.8 kb: subcellular localization classification (iLoc-lncRNA D1, iLoc-lncRNA D2 and lncATLAS-Localization), continuous cytoplasmic-to-nuclear RCI regression (lncATLAS-RCI), coding potential prediction (GENCODE-Coding, RNAErnie-lncRNA and RNAChallenge), official variation-record status annotation (Disease), multi-context expression capacity prediction (Expression) and RBP binding (POSTAR3-UPF1). The second tier evaluates four mechanistic structural and post-transcriptional regulatory assays: pairwise RNA contact prediction (mRNA-Structure and lncRNA-Structure), nucleotide-resolution DMS reactivity profiling (mRNA-DMS), 5^′^ UTR mean ribosome load regression (5UTR-MRL), and splice-site recognition across donor and acceptor junctions (RiNALMo-Splice). Together, these 15 datasets interrogate three distinct modelling granularities: transcript-level classification and regression, nucleotide-level chemical reactivity profiling, and nucleotide-pair contact recovery.

All 15 datasets derive their underlying sequences, annotations, and experimental measurements from public repositories, with our contributions centring on task formulation, data curation, and unified benchmarking (Extended Data Tables 1 and 4). Specifically, the suite comprises three curation groups: (1) five task datasets formulated and constructed in this study from primary public databases—mapping public CLIP-seq peaks to lncRNA exons to define transcript-level UPF1 binding (POSTAR3-UPF1), linking official LncBook 2.0 genes (represented by their longest GRCh38 GTF transcripts) to official variation-record status (Disease) or five-context binarized expression capacity labels (Expression), and matching gene–cell-line fractionation readouts to longest annotated lncRNA transcripts to construct thresholded compartment classification (lncATLAS-Localization) and continuous ratio regression (lncATLAS-RCI); (2) six datasets adapted from existing task resources with substantial reprocessing, including merging, deduplication, label conversion, length filtering and resplitting (iLoc-lncRNA D2, mRNA-Structure, mRNA-DMS, 5UTR-MRL, RNAErnie-lncRNA and RiNALMo-Splice); and (3) four datasets incorporated from preprocessed sequence–label pairs or established splits (iLoc-lncRNA D1, the expanded GENCODE/RefSeq GENCODE-Coding artifact, RNAChallenge and lncRNA-Structure).

Thirteen datasets serve as internal benchmarks for downstream adaptation, whereas RNAChallenge and lncRNA-Structure are held out as external generalization tests evaluated directly using checkpoints adapted on the human GENCODE-Coding and mRNA-Structure cohorts, respectively, without further tuning. To isolate intrinsic representation quality from downstream overfitting and ensure comparable evaluations among LncRFM and seven established RNA foundation models, all models were evaluated with frozen encoders on identical data splits and test record lists within each dataset (Extended Data Tables 1 and 2).

### 2.3 The LncRFM architecture and pretraining mechanisms

LncRFM is designed to internalize secondary-structure topologies during pretraining while operating strictly on raw primary sequence during downstream inference (Fig. 1b,c). The encoder architecture couples a multiscale depthwise-separable convolutional stem (parallel kernel widths of 3 and 7) with a hybrid bidirectional backbone comprising ten state-space blocks interleaved with two self-attention layers. The convolutional stem resolves variable-length local sequence motifs before token representations propagate through the bidirectional state-space backbone, which integrates long-range contexts across up to 2,050 encoded tokens with near-linear computational complexity.

During pretraining on LncCorpus, predicted base-pairing partners guide representation learning through two synergistic mechanisms: (1) structure-guided joint masking, which masks base-paired nucleotides simultaneously to force reconstruction of long-range structural couplings from surrounding context; and (2) pair-gated fusion, which dynamically weights forward and reverse state-space trajectories according to each nucleotide’s pairing state. At downstream adaptation and inference, LncRFM discards structural inputs entirely, replacing pair-gated fusion with parameter-free directional state averaging. Consequently, adapting LncRFM to downstream tasks requires no external RNA folding step and updates only a lightweight multi-branch bottleneck adapter and task head (representing just 1.7% of total parameters in the coding-potential configuration) atop the frozen encoder to extract pooled transcript-level representations, nucleotide-wise states or symmetric pairwise contact matrices across tasks (Fig. 1c).

### 2.4 Generalizable performance across functional and structural granularities

Evaluated across all 15 LncBench datasets under the frozen-encoder protocol, LncRFM achieved the highest primary-metric score on 11 of the 13 internally adapted tasks (Fig. 2a–m), while closely matching the top structure-specialized baseline on external human lncRNA contact prediction and showing moderate cross-kingdom transfer on external coding-potential classification (Fig. 2n,o). In the first tier of transcript-level lncRNA functional and annotation tasks, gains spanned binary compartment classification, multilabel expression and localization sorting, and continuous cytoplasmic-to-nuclear ratios. Relative to the strongest baseline on each task’s primary metric, LncRFM raised F1 on lncATLAS-Localization from 0.520 to 0.583 (0.58 versus 0.52 rounded; Fig. 2d), Pearson correlation *r* on lncATLAS-RCI from 0.344 to 0.403 (Δ*r* = 0.059 over LncRNA-BERT; Fig. 2h) and Micro-F1 on iLoc-lncRNA D1 from 0.743 to 0.764 (0.76 versus 0.74 rounded; Fig. 2f), while achieving comparable Micro-F1 on the heavily class-imbalanced 3.8-kb iLoc-lncRNA D2 dataset (0.769925 versus 0.766917 for LncRNA-BERT, both 0.77 rounded; Fig. 2g, analyzed further in Section 2.6). Consistent leads also emerged across RBP recognition, multi-context expression capacity profiling, official variation-record annotation and coding-potential discrimination: LncRFM surpassed the top baseline on each of these primary metrics—including POSTAR3-UPF1 (AUROC 0.757 versus 0.714 for structRFM and LncRNA-BERT, Δ = 0.043; Fig. 2a), Expression (Macro-AUPRC 0.498 versus 0.470 for LncRNA-BERT, Δ = 0.028; Fig. 2b), Disease (AUROC 0.791 versus 0.740 for LncRNA-BERT, Δ = 0.052; Fig. 2c) and GENCODE-Coding (AUROC 0.980 versus 0.968 for LncRNA-BERT, Δ = 0.012; Fig. 2e).

Notably, LncRFM’s advantage extended to the second tier of quantitative translation and nucleotide- or pair-resolution structural assays, where the structure-pretrained structRFM model consistently served as the strongest baseline. Operating directly on primary sequence without folding inputs, LncRFM increased squared Pearson correlation on 5UTR-MRL from 0.28 to 0.35 (Fig. 2k), base-pair contact F1 on mRNA-Structure from 0.12 to 0.16 (Fig. 2i) and nucleotide-level reactivity correlation on mRNA-DMS from 0.66 to 0.67 (Fig. 2j). On only two adapted tasks did alternative encoders lead: RNAErnie-lncRNA classification (AUROC 0.93 versus 0.94 for LncRNA-BERT; Fig. 2l) and short-window RiNALMo-Splice recognition (mean F1 0.64 versus 0.71 for RNA-FM; Fig. 2m). Finally, when the internally adapted checkpoints were transferred directly to external test sets without further tuning, LncRFM closely approached structRFM on human lncRNA-Structure (F1 0.140621 versus 0.141048, both 0.14 rounded; Fig. 2o), whereas on multi-species RNAChallenge it achieved a coding F1 of 0.37 (0.374 unrounded, ranking fourth behind RNA-FM at 0.43 [0.429]; Fig 2n).

### 2.5 Multi-metric robustness and scaling into the multi-kilobase regime

To verify whether benchmark-wide patterns held across evaluation criteria, we assessed 75 dataset–metric combinations across the 13 adapted datasets spanning threshold-dependent classification metrics (accuracy, balanced accuracy, precision, recall, specificity, F1 and MCC), threshold-free ranking and calibration metrics (AUROC, AP/AUPRC and Brier score), and regression correlation and error metrics (Pearson *r*, Spearman *ρ*, *R*^2^, MAE and RMSE) (Fig. 3a). Across LncBench, LncRFM led or tied for the highest unrounded score across all three micro-averaged metrics on both iLoc-lncRNA D1 and iLoc-lncRNA D2, all five regression metrics on lncATLAS-RCI (*r* = 0.402812, *ρ* = 0.383663, MAE = 1.261536, RMSE = 1.595131, *R*^2^ = 0.160802) and mRNA-DMS, and Pearson *r* and *r*^2^ on 5UTR-MRL. Among classification and structural tasks, LncRFM achieved the highest score on six of eight plotted metrics on lncATLAS-Localization (F1, AUROC, AUPRC, accuracy, precision and MCC) and POSTAR3-UPF1 (AUROC 0.757066, AP 0.701354, F1 0.673102, accuracy 0.670278, precision and MCC 0.366777, while ranking behind RNA-FM on recall and LncRNA-BERT on specificity), six of eight on Expression (Macro-AP 0.498310, Macro-AUROC 0.648401, Micro-AP 0.641916, Micro-AUROC 0.762081, Micro-F1 0.584510 and Hamming loss 0.283538, for which lower values indicate better performance, whereas LncRNA-BERT attained higher Macro-F1 of 0.330014 versus 0.321282 and exact match of 0.159141 versus 0.156691), five of eight on Disease (AUROC 0.791401, AP 0.810961, F1 0.753892, accuracy 0.717769 and MCC 0.435265, whereas LncRNA-BERT led in precision and specificity and RiNALMo micro led in recall), four of five on mRNA-Structure, and four of eight on GENCODE-Coding (AUROC 0.979960, AP 0.981557, precision 0.967863 and specificity 0.963809, whereas LncRNA-BERT achieved higher F1 of 0.941769 versus 0.929438 and accuracy of 0.935877 versus 0.925427, together with lower Brier error). Across all 75 dataset–metric combinations, LncRFM ranked first (including unrounded ties) in 49 combinations, achieving the highest mean relative score (0.976, ahead of 0.905 for structRFM and 0.873 for LncRNA-BERT), the highest median relative score (1.000, ahead of 0.960 for LncRNA-BERT and 0.945 for structRFM), and the narrowest interquartile range (IQR = 0.019, versus 0.087–0.181 across the seven baselines; Fig. 3b). Descriptively across the 13 primary metrics, LncRFM also achieved the best mean ordinal rank (1.462, compared with 2.923 for LncRNA-BERT and 3.308 for structRFM).

We next performed 2,000 paired record-level bootstrap resamples across ten core datasets to evaluate rank stability and primary-score margins relative to a fixed baseline comparator on each task (Fig. 3c). Across the test sets, primary-score differences over the fixed baseline were positive across all ten datasets, with 95% percentile intervals excluding zero on eight tasks (Fig. 3c): lncATLAS-Localization (ΔF1 = 0.0631; 95% CI, 0.0402–0.0866), lncATLAS-RCI (Δ*r* = 0.0592; 0.0427–0.0765), Disease (ΔAUROC = 0.0516; 0.0465–0.0569), POSTAR3-UPF1 (ΔAUROC = 0.0426 relative to structRFM; 0.0341–0.0505), mRNA-Structure (ΔF1 = 0.0352; 0.0321–0.0383), Expression (ΔMacro-AUPRC = 0.0284; 0.0218–0.0347), GENCODE-Coding (ΔAUROC = 0.0121; 0.0092–0.0151) and mRNA-DMS (Δ*r* = 0.0106; 0.0068–0.0144), whereas intervals crossed zero on iLoc-lncRNA D1 (ΔMicro-F1 = 0.0213; 0.0047–0.0489) and iLoc-lncRNA D2 (ΔMicro-F1 = 0.0030; 0.0131–0.0194).

Because attention-based encoders are restricted to 510–1,024 encoded tokens and must split longer transcripts across sliding windows, we examined performance across three transcript-length regimes (≤1, 024, 1, 025–2, 050 and *>* 2, 050 nt; Fig. 3d). Within the evaluated cohorts, LncRFM maintained high conditional bootstrap winning shares across length strata. On lncATLAS-Localization, rank-1 winning shares rose monotonically from 83.4% on transcripts ≤1, 024 nt to 93.4% between 1, 025 and 2, 050 nt, and 100% above 2, 050 nt, with the cytoplasmic F1 margin over the strongest baseline nearly tripling from 0.024 (≤1, 024 nt; F1 0.605 versus 0.580 for LncRNA-BERT) to 0.036 (1, 025–2, 050 nt; 0.525 versus 0.489 for RNA-FM) and 0.068 (*>* 2, 050 nt; 0.612 versus 0.543 for LncRNA-BERT). Across POSTAR3-UPF1, Expression, Disease, lncATLAS-RCI and GENCODE-Coding, LncRFM achieved near-complete rank-1 winning shares across all three length bins on Expression (100.00%, 100.00% and 100.00%), Disease (100.00%, 100.00% and 100.00%), GENCODE-Coding (100.00%, 99.85% and 100.00%) and lncATLAS-RCI (99.35%, 99.70% and 99.65%), and led all three strata on POSTAR3-UPF1 (100.00% at ≤ 1, 024 nt, 100.00% at 1, 025–2, 050 nt and 88.75% at *>* 2, 050 nt). Winning shares also remained high across longer regimes on iLoc-lncRNA D1 (92.6% at 1, 025–2, 050 nt and 95.5% above 2, 050 nt), reversed early deficits above 2, 050 nt on iLoc-lncRNA D2 (79.4%), and reached 100% in the sole populated length bin (≤1, 024 nt) for both mRNA-Structure and mRNA-DMS.

### 2.6 Calibration, biological gradients and fine-grained structural recovery

To understand how pretrained representations behave across distinct biological regimes, we examined joint metric trade-offs, probability calibration and quantitative biological gradients across tasks (Fig. 4). Across secondary metric profiles on 5UTR-MRL, RNAErnie-lncRNA and RiNALMo-Splice (Fig. 4a–c), LncRFM achieved the strongest correlation with experimental ribosome load on 5UTR-MRL (Pearson *r* = 0.595), whereas structRFM yielded a lower MAE (1.089 versus 1.144; Fig. 4a); on RNAErnie-lncRNA, LncRFM ranked second in both AUROC (0.932) and F1 (0.855) behind LncRNA-BERT (Fig. 4b); and for splice-site recognition on RiNALMo-Splice, LncRFM achieved higher F1 on acceptors than on donors (0.679 versus 0.604), ranking third and eighth among the eight encoders, respectively (Fig. 4c).

In binary classification tasks evaluated against LncRNA-BERT (the strongest baseline on each task’s primary metric), ranking, calibration and thresholded accuracy exhibited task-specific trade-offs (Fig. 4d). On POSTAR3-UPF1, Disease and lncATLAS-Localization, LncRFM simultaneously outperformed LncRNA-BERT in AUROC (0.757 versus 0.714, 0.791 versus 0.740, and 0.761 versus 0.692, respectively), average precision (0.701 versus 0.675, 0.811 versus 0.734, and 0.568 versus 0.460), Brier probability error (0.213 versus 0.218, 0.191 versus 0.209, and 0.207 versus 0.244) and F1 at the 0.5 cutoff (0.673 versus 0.633, 0.754 versus 0.676, and 0.583 versus 0.520). By contrast, on GENCODE-Coding, while LncRFM achieved higher threshold-free ranking scores than LncRNA-BERT (AUROC 0.980 versus 0.968 and AP 0.982 versus 0.965), LncRNA-BERT produced lower Brier error (0.054 versus 0.068) and higher F1 at the 0.5 threshold (0.942 versus 0.929). Across quantitative biological covariates (Extended Data Fig. 3a–c,e), predicted probabilities tracked empirical directional trends but displayed compressed dynamic ranges relative to observed frequencies: on GENCODE-Coding, mean predicted coding probabilities rose across longest-ORF coverage bins from 0.198 to 0.784 for LncRFM (versus 0.126 to 0.886 for LncRNA-BERT) compared with observed positive fractions from 0.104 to 0.925 (Extended Data Fig. 3a); on lncATLAS-Localization, LncRFM’s predicted cytoplasmic probabilities increased across experimentally measured cytoplasmic-to-nuclear log-ratios (Extended Data Fig. 3b); on POSTAR3-UPF1, predicted binding probabilities rose modestly across non-UPF1 RBP co-binding diversity bins (0.464 to 0.649 for LncRFM and 0.427 to 0.682 for LncRNA-BERT) compared with the steep observed gradient (0.139 to 0.962; Extended Data Fig. 3c); and on lncATLAS-RCI, predicted decile means tracked observed RCI ordering (1.561 to 0.652 for LncRFM) but exhibited substantial shrinkage toward the dataset mean relative to observed decile means (4.364 to 1.791; Extended Data Fig. 3e).

Across the 13 labels in the three multilabel datasets (Expression, iLoc-lncRNA D1 and iLoc-lncRNA D2), LncRFM’s label-wise F1 correlated strongly with label prevalence (Spearman *r_s_* = 0.96, unrounded 0.963882; Fig. 4e), reflecting high F1 on prevalent categories such as *normal tissue cell line* (0.782, prevalence 0.644) and *organ development* (0.615, prevalence 0.484) in Expression, alongside persistent difficulty on rare labels such as *cell differentiation* (0.018, prevalence 0.164) and *subcellular localization* (0.051, prevalence 0.172), as well as zero F1 on four rare localization classes in iLoc-lncRNA D1/D2. In multilabel prediction diagnostics (Fig. 4f), LncRFM exhibited the smallest magnitude of label-count under-prediction bias on Expression (0.19 versus 0.28 to 0.43 across baselines), with a dominant label-set share of 0.53 and an exact-match rate of 0.16 (close to 0.16 for LncRNA-BERT, or 0.156691 versus 0.159141 unrounded), while leading sample-wise F1 across transcripts with one to three true labels (Fig. 4g). On iLoc-lncRNA D1, LncRFM achieved the highest exact-match rate (0.42 versus 0.28–0.36 across baselines; dominant share 0.83), whereas on iLoc-lncRNA D2, LncRFM’s dominant-label share of 0.94 reflected predictions restricted solely to *Exosome* in 94% of records (exact match 0.77).

At base-pair and single-nucleotide resolution, transcript-level diagnostics on the Rouskinlab mRNA cohorts confirmed that structure-guided pretraining internalizes long-range pairing constraints into sequence representations. In mRNA-Structure con-tact prediction, improvements reflected simultaneous gains in precision and recall, with LncRFM exceeding structRFM on both measures for 61% of the 291 test transcripts and reaching a pooled precision of 0.095 and recall of 0.424. Crucially, mean transcript-level F1 differences over structRFM remained positive across all five groups stratified by the fraction of long-range base pairs (Extended Data Fig. 1b, c). For mRNA-DMS reactivity prediction, LncRFM achieved lower MAE in transcript groups with higher pairing density and lower mean reactivity—where base-paired stems shield nucleotides from chemical modification—whereas structRFM had lower MAE in several higher-reactivity groups (Extended Data Fig. 1d).

### 2.7 Out-of-distribution generalization across species and RNA biotypes

To test whether adapted representations generalize across evolutionary and biotype boundaries without further training, we evaluated coding-potential and RNA contact checkpoints adapted on the human GENCODE-Coding and mRNA-Structure internal cohorts directly on the external RNAChallenge and lncRNA-Structure benchmarks, respectively (Fig. 5). On RNAChallenge (*n* = 27, 283 records), which spans animal, plant and fungal transcripts beyond the human scope of internal training, cross-species domain shift reduced F1 across all eight encoders relative to the human internal coding cohort, with a median change of 0.451 (Fig. 5a). Under this shift and an overall score inversion across non-human clades that depressed global AUROC across all human-trained encoders, RNA-FM retained the highest external F1 (0.429) at the prespecified decision threshold, followed by RNAErnie (0.399), structRFM (0.396) and LncRFM (0.374, down from 0.929 internally), whereas UTR-LM led in external average precision (0.494), AUROC (0.335) and accuracy (0.372).

Pairwise contact prediction exhibited strong cross-biotype stability when transferring checkpoints trained on human mRNAs to external human lncRNAs (Fig. 5b and Extended Data Fig. 3h). Across the eight encoders, the median F1 change was only −0.003, and model rankings were strongly preserved between the internal mRNA and external lncRNA sets (Spearman *r_s_* = 0.90). On the lncRNA-Structure benchmark, comprising 30 structure records representing 24 unique sequences, LncRFM achieved a contact F1 of 0.141 (0.140621 unrounded, compared with 0.155 internally), closely matching structRFM (0.141048 unrounded; both 0.141 displayed) while ranking second in AP (0.101 versus 0.104) and first in AUROC (0.962; Fig. 5b).

### 2.8 Task-dependent contributions of local motif extraction and pairing-guided pretraining

To assess the contributions of the three structure-aware components, we compared the full LncRFM model with a base encoder lacking the motif stem, joint masking and pair-gated fusion, as well as three leave-one-out variants across the 13 adaptation tasks under matched pretraining and evaluation protocols (Fig. 6a; Extended Data Table 1). The full model outperformed the base encoder on 10 tasks, tied it on RNAErnie-lncRNA, and scored lower on lncATLAS-Localization and RiNALMo-Splice. It achieved the highest unrounded score among the five configurations on eight tasks. Representative full-versus-base gains included an AUROC increase from 0.765 to 0.791 on Disease, a Macro-AUPRC increase from 0.465 to 0.498 on Expression, a Micro-F1 increase from 0.715 to 0.770 on iLoc-lncRNA D2, and an increase in Pearson’s *r* from 0.387 to 0.403 on lncATLAS-RCI.

The leave-one-out comparisons revealed task-dependent effects (Fig. 6b). Pair-gated fusion produced the largest observed omission effect on seven tasks where the full model outperformed the base encoder, including Disease (ΔAUROC = 0.038), lncATLAS-RCI (Δ*r* = 0.032) and 5UTR-MRL (Δ*r*^2^ = 0.032). Joint masking produced the largest observed omission effect on iLoc-lncRNA D1 (ΔMicro-F1 = 0.062) and lncATLAS-Localization (ΔF1 = 0.034), whereas the motif stem produced the largest observed omission effect on Expression (ΔMacro-AUPRC = 0.029) and mRNA-DMS (Δ*r* = 0.004).

The benefits were not uniform across tasks. The motif-free variant achieved the highest score on RNAErnie-lncRNA, lncATLAS-Localization, mRNA-Structure and RiNALMo-Splice, whereas the variant without joint masking was marginally highest on 5UTR-MRL.

## 3 Discussion

Decoding the functional landscape of long non-coding RNAs from primary sequence requires reconciling two distinct spatial scales: short nucleotide motifs that mediate local protein or ribosome recognition, and kilobase-spanning secondary structures that govern global transcript architecture and compartment sorting. By coupling a multiscale convolutional motif stem with a hybrid bidirectional state-space backbone pretrained on the 24.7-million-sequence LncCorpus resource, LncRFM processes transcripts of up to 2,050 tokens within a single context window while handling even longer transcripts through overlapping windows. Across LncBench, LncRFM achieved strong rank stability into the multi-kilobase regime (*>* 2, 050 nt) and substantial gains in threshold-free ranking on tasks such as nuclear-versus-cytoplasmic localization (AUROC rising from 0.692 to 0.761 on lncATLAS-Localization) and variation-record status classification (AUROC rising from 0.740 to 0.791 on Disease), translating directly into more reliable candidate prioritization across transcriptome-wide screens.

Evaluating representation learning across this length and resolution spectrum required a dedicated multi-scale benchmark. LncBench unifies kilobase-scale lncRNA functional and annotation readouts with nucleotide- and base-pair-resolution structural and regulatory assays under a strictly controlled frozen-encoder protocol. Rather than relying solely on point-estimate accuracy, LncBench incorporates length-stratified bootstrap resampling, probability calibration, multilabel mode-collapse diagnostics and quantitative biological gradient tracking. Specifically, while LncRFM predictions align monotonically with biological covariates such as cytoplasmic-to-nuclear localization ratios, open-reading-frame coverage, RCI deciles and long-range base-pairing burdens (≥128 nt), predicted probabilities and continuous RCI estimates exhibit com-pression toward dataset means, and rare multilabel categories remain challenging to recover at fixed decision thresholds. Furthermore, although LncRFM led threshold-free ranking (AUROC and AP) on GENCODE-Coding, LncRNA-BERT achieved lower Brier error and higher F1 at the 0.5 cutoff, underscoring that ranking superiority does not automatically confer optimal thresholded calibration across all tasks.

A central design principle of LncRFM is confining structural supervision to pre-training while preserving sequence-only downstream inference. During pretraining on LncCorpus, predicted base-pairing partners link reconstruction targets across distal stem regions via joint masking, while binary pairing status modulates directional state integration through pair-gated fusion. Because downstream adaptation replaces pair-gated fusion with parameter-free directional state averaging, structural priors from 24.7 million transcripts are distilled into the sequence encoder itself, eliminating the computational bottleneck and prediction noise of running external folding algorithms on every new transcript at inference. The ablations indicate partially overlapping and task-dependent contributions rather than a universal mechanistic division of labour. Pair-gated fusion showed the largest observed omission effect on several interaction, localization, coding and structural tasks; joint masking showed the largest effect on iLoc-lncRNA D1 and lncATLAS-Localization; and the motif stem showed the largest effect on Expression and mRNA-DMS. The higher scores of compact variants on localization and splice prediction further show that the complete architecture is not uniformly optimal across downstream targets.

Our out-of-distribution evaluations further highlight distinct generalization regimes across biotype and evolutionary boundaries. Pairwise contact-prediction rankings transferred seamlessly from internal human mRNAs to external human lncRNAs (Spearman *r_s_* = 0.90), indicating that base-pairing grammar learned by LncRFM generalizes across RNA biotypes. By contrast, transferring human-adapted coding-potential checkpoints to the multi-kingdom RNAChallenge collection spanning animal, plant and fungal transcripts induced a pronounced domain shift across all eight foundation models (median ΔF1 = 0.451), reducing LncRFM’s F1 from 0.929 to 0.374. Including both internal adaptation splits and held-out external benchmarks within LncBench exposes these transfer boundaries alongside standard within-distribution comparisons.

Further work can extend LncRFM’s adaptation strategies, structural supervision and species coverage. First, to isolate intrinsic representation quality across all eight foundation models, our benchmark enforced a frozen-encoder protocol with a lightweight bottleneck adapter (1.7% trainable parameters in the binary reference configuration); exploring full-encoder fine-tuning or low-rank adaptation across LncBench may unlock additional task-specific capacity. Second, the base-pairing topologies in LncCorpus were derived computationally via ensemble folding and LinearFold rather than experimental probing; integrating high-throughput in vivo chemical probing data (such as DMS-MaPseq or SHAPE-MaP profiles) into pretraining could provide higher-fidelity structural supervision and capture dynamic cellular conformations. Third, while LncBench centres primarily on human lncRNA functional assays, expanding multi-task annotations to non-human vertebrates, plants and microbes—and distinguishing official database annotation status (such as variation-record presence in Disease) from experimentally validated causal mechanisms—will be essential for establishing pan-eukaryotic non-coding RNA foundation models.

In summary, LncCorpus, LncBench and LncRFM establish an open data, benchmarking and modelling ecosystem for kilobase-scale RNA representation learning. Achieving top primary-metric performance on 11 of 13 adapted tasks and strong cross-biotype generalization on external lncRNA structure prediction, LncRFM demonstrates that pairing-guided state-space pretraining effectively bridges local sequence grammar and global structural context. More broadly, sequence-only encoders inter-nalized with structural priors offer a scalable foundation for downstream molecular engineering, including programmatic nucleic-acid aptamer design [40], therapeutic RNA optimization and de novo antibody design [41].

## 4 Methods

### 4.1 LncCorpus and LncBench curation

#### LncCorpus assembly and filtering

To construct LncCorpus, we combined RNAcentral transcript sequences [42] with secondary-structure pairing indices derived either from the multimethod MUSES ensemble used by structRFM (≤512 nt; approximately 21.5 million sequences) or from LinearFold [17, 39] for longer transcripts. Specifically, we contributed over 9.6 million additional long noncoding and transcript sequences to assemble an initial source corpus of 31,121,367 records. Filtering this source corpus to sequences between 200 and 2,050 nt with strictly aligned sequence and pairing-vector lengths excluded transcripts shorter than 200 nt and retained 24,771,981 high-quality records (15.1 million from the MUSES subset and 9.6 million long sequences), comprising 24,648,121 sequences for model pretraining and 123,860 sequences for a held-out validation partition. Spanning 24.6 million training records across kilobase-scale lncRNAs, mRNAs and other non-coding RNA classes, LncCorpus is among the largest structure-annotated corpora compiled for RNA foundation model pretraining. Detailed provenance and step-by-step filtering counts are documented in Extended Data Tables 4 and 5.

#### LncBench task collection and provenance tiers

LncBench comprises 15 downstream datasets spanning 10 prediction tasks across three modelling granularities: transcript-level classification and regression, nucleotide-level reactivity regression, and nucleotide-pair contact prediction. Thirteen datasets provide training, validation and test partitions for downstream adaptation, whereas RNAChallenge and lncRNA-Structure serve as external generalization benchmarks evaluated directly using internally adapted checkpoints without further tuning. All 15 datasets derive their underlying sequences, annotations and experimental measurements from public repositories, with our contributions centring on task formulation, data standardization and unified benchmarking (Extended Data Tables 1 and 4). Grouped by curation workflow and provenance, the 15 datasets comprise: (1) five task datasets newly formulated and constructed in this study from primary public databases (POSTAR3-UPF1, Disease, Expression, lncATLAS-Localization and lncATLAS-RCI); (2) six datasets adapted from existing task resources with substantial in-house reprocessing, including multi-source merging, deduplication, label conversion, length filtering and re-splitting (iLoc-lncRNA D2, mRNA-Structure, mRNA-DMS, 5UTR-MRL, RNAErnie-lncRNA and RiNALMo-Splice); and (3) four datasets incorporated from preprocessed sequence–label pairs or established data splits (iLoc-lncRNA D1, GENCODE-Coding, RNAChallenge and lncRNA-Structure).

#### Subcellular localization and RCI regression

iLoc-lncRNA D1 interfaces directly with the compiled iLoc-lncRNA sequence–compartment table [43], whereas iLoc-lncRNA D2 was reprocessed from human lncRNA records in NCRNALocate-EL [44]. Across both datasets, transcript identifiers and compartment ontology terms were harmonized, and all compartments assigned to a transcript were encoded as a multihot target vector. For lncATLAS-Localization and lncATLAS-RCI, we curated gene–cell-line nuclear and cytoplasmic fractionation measurements from lncATLAS [36], normalized Ensembl gene identifiers and linked each gene to its longest annotated lncRNA transcript in GENCODE v47 [45]. Each record retains its gene–cell-line measurement rather than collapsing all cell lines into a single value per gene. The processed metric ratio2 represents the cytoplasmic-to-nuclear relative concentration index (RCI), defined as log_2_(cytosol*/*nucleus). In lncATLAS-Localization, records with RCI ≥ 1 (cytoplasmic-enriched) and RCI 1 (nuclear-enriched) were assigned binary labels of 1 and 0, respectively, excluding intermediate values; lncATLAS-RCI retains the continuous RCI value (17, 083 training, 3, 727 validation and 3, 411 test records) as a regression target.

#### Coding potential

GENCODE-Coding comprises a curated collection of human lncRNA and protein-coding transcripts compiled from GENCODE [45] and RefSeq [46] annotations (131, 264 training, 7, 271 validation and 6, 316 test transcript records). RNAChallenge was incorporated in its established format to provide an external multispecies evaluation of binary coding-potential classification using the checkpoint adapted on the human GENCODE-Coding splits.

#### Disease-association and expression-context annotations

Disease and Expression were formulated from official LncBook 2.0 human lncRNA gene tables [37], with each gene represented by its longest transcript sequence in the official GRCh38 GTF (95, 243 prepared gene records corresponding to 94, 291 exact-unique sequences and 91, 564 clusters at 90% sequence identity). Disease defines a binary classification target indicating whether a gene has any recorded annotation in the official LncBook variation table (aggregating entries from dbSNP, COSMIC [47], ClinVar [48] and GWAS); thus, positive labels denote the presence of an official variation record rather than confirmed disease causality or exclusively pathogenic variants. Expression consolidates five official expression-capacity contexts (*normal tissue cell line*, *organ development*, *preimplantation embryo*, *cell differentiation* and *subcellular localization*) into a five-dimensional binary multilabel target by mapping high, medium or low expression capacity (HC, MC or LC) to 1 and no expression (NE) to 0. Detailed target construction rules and label encodings are provided in Supplementary Methods, Section 2.4.

#### RNA–protein binding

POSTAR3-UPF1 was constructed by intersecting human CLIP-seq binding peaks from POSTAR3 [35] with GENCODE v47 lncRNA exon coordinates matched by chromosome, strand and genomic interval, after excluding generic RBP occupancy entries. A transcript was labelled positive if at least one mapped peak supported the interaction. Among the RBPs catalogued in POSTAR3, UPF1 was selected as the primary benchmark target for transcript-level lncRNA–RBP interaction prediction (43, 078 training, 8, 876 validation and 8, 216 test transcripts).

#### RNA contact and DMS reactivity profiling

The reprocessed Rouskinlab resource [38] comprises 1,397 human mRNAs with matched primary sequences, secondary structures and in vivo dimethyl sulfate (DMS) reactivity measurements. From this resource, mRNA-Structure and mRNA-DMS share cluster-aware training and validation partitions of 1,019 and 146 transcripts, respectively, with scored test sets comprising 291 transcripts for mRNA-Structure and 232 for mRNA-DMS (Extended Data Table 1). In mRNA-Structure, secondary structures were converted into undirected nucleotide-pair contact sets. In mRNA-DMS, the normalized reactivity at each assayed nucleotide serves as a continuous regression target; valid reactivities lie in [0, 1], whereas sentinel values (999) were masked as miss-ing and excluded from loss and metric computations. Thirty human lncRNA structure records processed in the identical format constitute the external lncRNA-Structure benchmark (representing 24 unique sequences).

#### Mean ribosome load, secondary coding classification and splice-site recognition

5UTR-MRL, RNAErnie-lncRNA and RiNALMo-Splice were adapted and standardized from the processed downstream benchmark collection of ref. [49]. These datasets evaluate 5^′^ UTR mean ribosome load (MRL) regression, binary lncRNA-versus-mRNA coding classification, and splice-junction recognition across 400-nt sequence windows, respectively. RiNALMo-Splice trains separate binary classifiers for donor and acceptor sites, using the unweighted mean of donor and acceptor F1 scores as the primary task metric.

#### Partitioning, redundancy control and preprocessing

To ensure comparable evaluations across all eight foundation models, all methods were trained and evaluated on identical training, validation and test partitions within every task (for example, 648*/*81*/*72 records for iLoc-lncRNA D1, 3, 170*/*396*/*317 for iLoc-lncRNA D2, 17, 083*/*3, 727*/*3, 411 for lncATLAS-RCI, and 232 test transcripts for mRNA-DMS; Extended Data Table 1). Although Expression and Disease have identical partition sizes (76, 196*/*9, 524*/*8, 571), they do not share an identical test gene set; within each task, all eight models were evaluated on the exact same test records. Sequence redundancy at 90% identity was controlled via MMseqs Linclust followed by exact and reverse-complement sequence cluster merging for Expression and Disease (91, 564 clusters), and via CD-HIT [50] for the remaining clustered datasets (40, 599 clusters for POSTAR3-UPF1 and 5, 871 for lncATLAS-RCI; Extended Data Fig. 2 and Extended Data Table 5). Exact cross-split sequence overlap among the training, validation and test partitions was verified to be strictly zero across POSTAR3-UPF1, Expression, Disease, lncATLAS-RCI and GENCODE-Coding. The two external generalization benchmarks (RNAChallenge and lncRNA-Structure) were evaluated in full using fixed checkpoints from the internal coding and structure runs. Each model processed IUPAC ambiguity codes using its native tokenizer; LncRFM includes a dedicated N token and maps other non-canonical symbols to [UNK]. Continuous targets for RCI and MRL were retained on their native scales.

### 4.2 LncRFM architecture and structure-guided pretraining

LncRFM combines a multiscale convolutional motif stem with a 12-block hybrid bidi-rectional encoder comprising ten selective state-space blocks and two self-attention blocks [34]. Predicted base-pairing topologies guide self-supervised pretraining through structure-guided joint masking and pair-gated bidirectional state fusion, whereas downstream inference operates on primary sequence alone. The pretrained encoder contains 48.10 million parameters, increasing to 48.96 million when paired with the downstream bottleneck adapter and binary classification head.

#### Input representation and tokenization

We stripped sequence whitespace, converted characters to uppercase and standardized thymine (T) to uracil (U). The single-nucleotide vocabulary is augmented with five special tokens: [CLS], [SEP], [PAD], [MASK] and [UNK]. The encoder supports a maximum context window of *L* = 2, 050 tokens with an embedding dimension of *d* = 768. Sequences exceeding the maximum window length were partitioned into overlapping sliding windows during encoding, with token representations in overlapping regions averaged across windows to reconstruct full-length transcript states. The encoder omits learned absolute positional embeddings; instead, local depthwise convolutions, bidirectional state-space scans and interleaved self-attention layers provide relative and global positional context.

#### Multiscale local motif stem

Let **X** R*^L^*^×*d*^ (*d* = 768) denote the input token embedding matrix. The motif stem applies two parallel one-dimensional depthwise-separable convolution branches with kernel widths of 3 and 7. Each branch performs a depthwise temporal convolution followed by a 1 × 1 pointwise linear projection and a GELU activation. The two branch outputs are concatenated along the feature dimension to form a 2*d*-dimensional representation, projected back to *d* dimensions via **W***_m_* ∈ R^2*d*×*d*^, and added to the input via dropout and a learnable scalar residual gate *α* ∈ R:

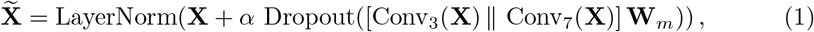

Where||denotes channel-wise concatenation. We set the dropout rate to 0.1 and ini-tialized *α* = 0 so that the motif branch contributed zero signal at initialization and gradually learned local motif representations during pretraining.

#### Bidirectional state-space and attention mixing

The encoder backbone consists of 12 residual blocks: blocks 6 and 12 use multi-head self-attention with 12 heads (head dimension 64), whereas the remaining ten blocks use non-causal bidirectional selective state-space mixers [34] configured with state dimension 64, local convolution width 7, expansion factor 2 and head dimension 64. Each state-space mixer scans the valid (non-padded) token sequence in both forward (5^′^ → 3^′^) and reverse (3^′^ → 5^′^) orientations and inverts the reverse output back to the native sequence order before directional fusion.

#### Pair-gated bidirectional fusion

During pretraining, each nucleotide *i* is accompanied by a binary pairing indicator *m_i_* ∈ {0, 1}, where *m_i_* = 1 if nucleotide *i* has a predicted base-pairing partner and *m_i_* = 0 otherwise. Each state-space block *k* computes a position-wise scalar gate with learnable parameters *w_k_, b_k_* ∈ R:

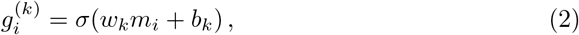

where *σ*(·) is the logistic sigmoid function, and combines the forward state h⟶^(k)^_i_ ∈ R*^d^* and reverse state h⟵^(k)^_i_ ∈ R*^d^* as

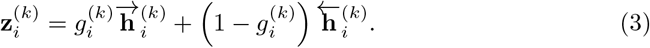

We initialized *w_k_* = *b_k_* = 0, assigning equal weights (*g*^(*k*)^ = 0.5) to both directions at the start of training. While the fusion gate receives only binary pairing status *m_i_*, the explicit partner index determines joint masking during pretraining. At downstream adaptation and inference, directional states are combined by parameter-free arithmetic averaging (h^(k)^_i_ = ½[h⟶^(k)^_i_ + h⟵^(k)^_i_] from primary sequence alone.

#### Masked RNA modelling

We pretrained the encoder from random initialization to reconstruct nucleotide identities from corrupted inputs. Let **X**^^^ _\M_ denote the corrupted sequence representation and M the final set of supervised positions. The cross-entropy reconstruction loss was

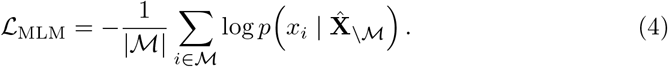

Under standard independent masking, non-special tokens were sampled independently with probability 0.15; of these, 80% were replaced by [MASK], 10% by a random nucleotide token and 10% left unchanged [51]. Nucleotide reconstruction served as the sole pretraining objective.

#### Structure-guided joint masking of pairing partners

To prevent local interpolation across base-paired stems, joint masking was introduced after an initial independent-masking warm-in phase covering approximately 42% of pretraining. Let *b* denote the cumulative number of batches processed by the data collator during the 300,000-update run, with two microbatches per optimizer update and approximately *B*_run_ = 6.0 × 10^5^ collator batches in total. After *b*_0_ ≈ 2.52 × 10^5^ batches, the probability *q*(*b*) of selecting joint masking for a batch increased linearly to *q*_end_ ≈ 0.148 at the end of the run:

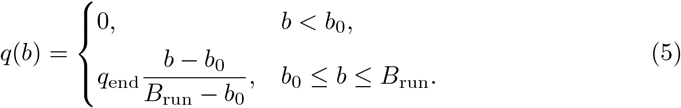

When joint masking was activated for a batch, non-special tokens were first sampled at the 0.15 base rate, and the predicted pairing partner of each selected paired nucleotide was simultaneously added to *M* (undergoing the same 80%/10%/10% token corruption rule) so that both paired nucleotides contributed jointly to the reconstruction loss. Unpaired nucleotides retained standard masking. Averaged over the entire run, this curriculum yields an expected joint-masking fraction of approximately 4% of collator batches. Binary pairing indicators *m_i_* were supplied to the pair-gated fusion layers throughout all 300,000 updates, including the initial independent-masking phase.

#### Pretraining optimization

Pretraining used four data-parallel GPU processes in FP32 precision. With a per-process microbatch size of 20 and gradient accumulation of 2, the effective global batch size was 160 sequences per optimizer update (corresponding to approximately two epochs across the 24.6-million-sequence training corpus over 300,000 updates). We used AdamW with a peak learning rate of 2 10^−4^, *β*_1_ = 0.9, *β*_2_ = 0.999, *ɛ* = 10^−8^ and weight decay 0. The learning rate warmed up linearly over the first 1,000 updates and decayed linearly thereafter. Global gradient norms were clipped at 0.5, and batches were grouped by sequence length to minimize padding. All downstream experiments initialized the encoder from the final 300,000-step checkpoint.

### 4.3 Downstream adaptation and evaluation protocol

#### Unified multi-branch bottleneck adapter

For each dataset requiring downstream adaptation, we initialized the encoder from its pretrained checkpoint, froze all encoder parameters and updated only a lightweight adapter module and prediction head [52]. Given frozen token representations, the adapter applies LayerNorm and projects token states to a 192-dimensional bottleneck. Bottleneck states pass in parallel through local one-dimensional convolutional branches with kernel widths of 3, 5 and 7, dilated width-3 convolutional branches with dilation rates of 2 and 4, and a global-context branch derived from the masked sequence mean. Softmax routing weights coupled with a position-wise sigmoid gate combine the six branches, and the aggregated output is projected back to the native encoder dimension and added to the input via a residual connection. Transcript-level tasks apply masked mean pooling over valid nucleotide states, whereas nucleotide- and pair-level tasks retain position-level states **h***_i_ ∈*R*^d^*. In the LncRFM binary classification reference configuration, 0.825 million parameters (0.825093 M) were trainable, corresponding to 1.7% of the 48.960 M total parameters (varying across task heads, such as 0.826121 M trainable of 48.961016 M total for five-label Expression and 1.219333 M trainable of 49.354228 M total for lncATLAS-RCI regression; Extended Data Table 2).

#### Prediction heads and loss functions

For binary and multilabel classification tasks, the pooled transcript representation passes through a two-layer MLP (hidden dimension 256) to produce class logits *y*^*_c_*. Binary tasks minimize class-weighted binary cross-entropy (BCE): given positive and negative training counts *n*_+_ and *n*_−_, initial inverse-frequency weights 1*/n*_+_ and 1*/n*_−_ are rescaled to yield a mean sample weight of 1 across the training split. Multilabel tasks use unweighted BCE (*w_yc_* = 1). Across *C* outputs, the unified loss is

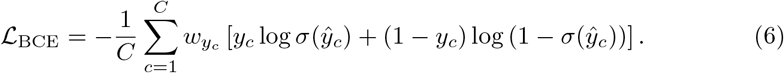

Except for pairwise contact prediction, all binary and multilabel predictions use the prespecified probability threshold of 0.5.

For continuous regression tasks, lncATLAS-RCI and 5UTR-MRL use transcript-level scalar MLPs, and mRNA-DMS uses a nucleotide-level MLP. All three minimize mean squared error (MSE),

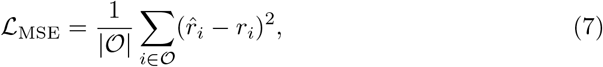

Where *O* indexes transcripts or nucleotide positions with valid observed targets (excluding special tokens and missing DMS positions).

For pairwise contact prediction (mRNA-Structure and external lncRNA-Structure), adapted nucleotide states **h***_i_* ∈ R*^d^* are projected to 256-dimensional query and key vectors via **W***_q_,* **W***_k_* ∈ R^256×*d*^ and combined with position-wise scalar unary terms *u_i_, u_j_* ∈ R (produced by a linear projection of **h***_i_* and **h***_j_*) to yield symmetric contact scores *s*∼*_ij_*:

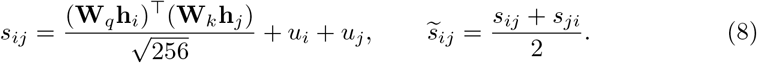

Contact metrics were computed over candidate canonical (A–U, U–A, C–G, G–C) and wobble (G–U, U–G) base pairs separated by more than three nucleotides (|*i-j|>* 3), augmented with all valid reference-positive contacts so that non-canonical or short-range true contacts were retained in the evaluation domain. The identical candidate rule was applied to all eight models within their effective input ranges. Training used positively weighted BCE with the positive-class weight set to the training ratio of candidate non-contacts to true contacts, capped at 200. The probability threshold was selected by grid search over {0.05, 0.10*, …,* 0.95} to maximize validation F1 on mRNA-Structure and held fixed for both the internal mRNA-Structure test set and the external lncRNA-Structure benchmark.

#### External generalization evaluation

RNAChallenge was evaluated using the checkpoint selected on the human GENCODE-Coding cohort, and lncRNA-Structure using the checkpoint selected on mRNA-Structure. Model parameters, prediction thresholds and checkpoint selections were frozen at their internally selected values across all eight methods.

### 4.4 Experimental settings and statistical analysis

#### Downstream training settings

Downstream experiments were implemented in PyTorch [53] and optimized with AdamW [54] using an effective batch size of 32, gradient-norm clipping at 1.0 and random seed 42 (Extended Data Table 3). For POSTAR3-UPF1, Expression, Disease, lncATLAS-RCI and GENCODE-Coding, LncRFM accumulated gradients to an effective batch size of 32 using microbatch size 16 with 2 accumulation steps for POSTAR3-UPF1, Expression, Disease and lncATLAS-RCI, and microbatch size 4 with 8 accumulation steps for GENCODE-Coding. Maximum training epochs were set to 100 for POSTAR3-UPF1, Expression, Disease, iLoc-lncRNA D1, iLoc-lncRNA D2 and lncATLAS-RCI; 60 for lncATLAS-Localization; 40 for GENCODE-Coding, mRNA-Structure, mRNA-DMS, 5UTR-MRL and RNAErnie-lncRNA; and 5 for each RiNALMo-Splice classifier. For validation-based checkpoint selection, large-scale transcript datasets (POSTAR3-UPF1, Expression, Disease, lncATLAS-RCI and GENCODE-Coding) used an early-stopping patience of 5 epochs with a minimum validation improvement of 0.001, whereas smaller or multi-stage adaptation tasks used a 15-epoch warmup window and a patience of 10 epochs (except the 5-epoch splice classifiers). Across all tasks, test evaluations were performed using the checkpoint selected by the validation criterion summarized in Extended Data Table 3. For LncRFM, dropout (0.1) and weight decay (0.01) were applied in Expression, iLoc-lncRNA D1, iLoc-lncRNA D2, lncATLAS-RCI, mRNA-Structure, mRNA-DMS and 5UTR-MRL, and set to 0 in POSTAR3-UPF1, Disease, GENCODE-Coding and the remaining binary classification tasks. Task-specific hyperparameters and early-stopping criteria are summarized in Extended Data Table 3.

#### Comparison models and input ranges

The seven baseline foundation models were structRFM, LncRNA-BERT, RNA-FM, SpliceBERT, RNAErnie, UTR-LM and RiNALMo micro [9–14, 17]. Each model used its officially released pretrained weights and native tokenizer (non-overlapping 3-mers for LncRNA-BERT; single nucleotides for all others). Configured maximum encoded input lengths in these experiments were 2,050 tokens for LncRFM; 1,024 for RNA-FM, UTR-LM and RiNALMo micro; 768 encoded 3-mer tokens for LncRNA-BERT; 512 for structRFM and RNAErnie; and 510 for SpliceBERT (Extended Data Table 2). For LncRFM, transcripts exceeding 2,050 tokens were processed using overlapping sliding windows (overlap 256 tokens) with overlapping token representations averaged prior to task-head pooling or pairwise scoring; baseline models processed long sequences according to their task run configurations (overlapping windows or truncation where configured in the baseline run logs). All eight encoders remained frozen and shared the same adapter architecture, task head design and data partitions within each task.

#### Ablation experiments

We compared the full LncRFM encoder with a base encoder lacking the motif stem, pairing-partner joint masking and pair-gated fusion, together with three leave-one-out variants that removed one component at a time. All variants were evaluated on the identical training, validation and test splits used in the main benchmark comparisons (Extended Data Table 1).

#### Evaluation metrics

Primary metrics were AUROC for POSTAR3-UPF1, Disease, GENCODE-Coding and RNAErnie-lncRNA; Macro-AUPRC (macro-averaged per-label average precision) for Expression; F1 for lncATLAS-Localization; Micro-F1 for iLoc-lncRNA D1 and D2; Pearson *r* for lncATLAS-RCI and mRNA-DMS; squared Pearson correlation (*r*^2^) for 5UTR-MRL; contact F1 for mRNA-Structure and lncRNA-Structure; and the unweighted mean of donor and acceptor F1 for RiNALMo-Splice. Secondary metrics included AUROC, average precision (AP/AUPRC), accuracy, balanced accuracy, precision, recall, specificity, Matthews correlation coefficient (MCC), Micro-Precision, Micro-Recall, Macro-F1, exact-match rate, Hamming loss, coefficient of determination (*R*^2^, computed as r2 score), mean absolute error (MAE), root mean squared error (RMSE) and the Brier probability error score [55]. Except for pairwise contact prediction, all binary and multilabel predictions used the prespecified probability threshold of 0.5.

Figure 3a,b summarizes 75 dataset–metric combinations across the 13 adapted datasets. To orient all metrics so that higher values consistently indicate better performance in visual summaries, error metrics *e* ∈ {Hamming loss, MAE, RMSE, loss} were mapped to 1*/*(1 + *e*) and MCC was linearly rescaled from [-1, 1] to [0, 1] via (MCC + 1)*/*2. Each unrounded transformed value was then divided by the unrounded maximum score achieved on that metric across the eight methods (assigning the top model a relative score of 1.000, with ties determined within a numerical tolerance of 10^−12^), while ordinal model rankings were computed from the original unrounded scores.

#### Rank tests and uncertainty analysis

We summarized descriptive primary-metric ordinal rankings across the 13 adapted datasets, assigning average ranks to ties.

For ten core datasets with record-level predictions, we performed 2,000 paired nonparametric bootstrap replicates [56] across test records (not independent model retrainings). Each replicate resampled test records with replacement using identical sample indices across all eight methods, recomputing primary metrics and model ranks from scratch. For Fig. 3c, the comparator on each task was a baseline held fixed across bootstrap replicates (structRFM on POSTAR3-UPF1, and the strongest test-set baseline on each of the other nine tasks, such as LncRNA-BERT on Expression, Disease, lncATLAS-RCI and GENCODE-Coding). Empirical 95% confidence intervals were defined by the 2.5th and 97.5th percentiles of the paired score differences.

For length-stratified analyses (Fig. 3d), test records were grouped by native sequence length into three regimes: 1, 024, 1,025–2,050 and *>* 2, 050 nt. Within each dataset–length bin, we performed 2,000 paired bootstrap resamples using identical sampled records across all eight methods. If *K* methods tied for the highest primary score in a replicate (within 10^−12^), each received a rank-1 contribution of 1*/K*, and the remaining methods received zero; the bootstrap rank-1 winning share was the mean contribution across all 2,000 resamples (reported as N/A for empty bins).

#### Prediction diagnostics and external-set uncertainty

Diagnostic analyses evaluated identical test records for paired model comparisons. Empirical quantiles defined six longest-ORF-fraction bins (*n* = 6, 316 on GENCODE-Coding), ten lncATLAS RCI deciles (*n* = 3, 411 on lncATLAS-RCI), five long-range-pair-fraction bins (*n* = 291 on mRNA-Structure) and six non-UPF1 RBP-diversity bins (*n* = 8, 216 on POSTAR3-UPF1), merging duplicate quantile boundaries where applicable. DMS reactivity diagnostics jointly stratified transcripts by quartiles of paired-nucleotide fraction and mean observed reactivity (*n* = 232). Wilson 95% confidence intervals were used for observed binary proportions, whereas mean predicted probabilities and paired transcript-level differences (LncRFM structRFM, such that negative ΔMAE indicates lower error for LncRFM) were estimated using 2,000 within-bin paired bootstrap replicates; for RCI deciles (Extended Data Fig. 3e), vertical bars depict prediction interquartile ranges (IQRs) rather than 95% confidence intervals. In multilabel diagnostics (Fig. 4e–g), dominant predicted label-set fraction (Dom.) denotes the proportion of records assigned the single most frequent predicted label combination, label-count bias denotes mean predicted minus mean observed positive labels per record, and sample-wise F1 was set to 0 when both true and predicted label sets were empty. For the 30 external lncRNA-Structure records, we stored transcript-level true positives, false positives and false negatives, resampled transcripts with replacement 2,000 times to recompute pooled precision, recall and F1, and quantified internal-to-external F1 rank preservation across methods using Spearman’s rank correlation (*r_s_*).

#### Software and reproducibility

All training and evaluation pipelines were executed using Python 3.9.23, PyTorch 2.3.1, HuggingFace Transformers 4.44.0, Datasets 3.1.0, NumPy 2.0.2, SciPy 1.13.1, scikit-learn 1.6.1 and pandas 2.3.3. Dataset construction rules, partition sizes, label distributions, sequence-overlap audits and file checksums are documented in the supplementary materials alongside Source Data for each figure panel.

## Data availability

Source Data summaries for the main figures, Extended Data Figs. 1–3 and Extended Data Tables 1–5 accompany the manuscript, including panel-level values, dataset-construction summaries and partition counts. The LncCorpus pretraining dataset [57] and the training, validation and test partitions of LncBench [58] are publicly avail-able on Zenodo. The source resources are available at: GENCODE [45]: https://www.gencodegenes.org/human/; RefSeq [46]: https://www.ncbi.nlm.nih.gov/refseq/; iLoc-lnc [43]: https://github.com/Superzchen/iLearnPlus; NCRNALocate-EL [44]: https://github.com/ze-liu/NCRNALocate-EL; lncATLAS [36]: https://lncatlas.crg.eu/; Lnc-Book [37]: https://ngdc.cncb.ac.cn/lncbook/; COSMIC [47]: https://cancer.sanger.ac.uk/cosmic; ClinVar [48]: https://www.ncbi.nlm.nih.gov/clinvar/; POSTAR3 [35]: https://postar.ncrnalab.org/; Rouskinlab [38]: https://github.com/rouskinlab; CD-HIT [50]: https://github.com/weizhongli/cdhit.

## Code availability

Code for LncRFM pretraining, pretrained weights, downstream fine-tuning and evaluation is available at https://github.com/chuyhu/LncRFM.

## Ethics declarations

The study used public RNA sequences, annotations and structure-probing data. It involved no new participant recruitment, tissue collection, animal experiments or clinical intervention; ethics approval was therefore not required.

## Author contributions

S.K.Z. and P.X. conceived and supervised the study. H.Z. conceptualized the idea, co-supervised the study, contributed data collection and framework implementation. A.C. designed and implemented LncRFM, curated the LncBench datasets, ran pretraining and downstream evaluations, analyzed the results and drafted the manuscript. R.L. contributed to benchmark evaluation and manuscript revision. H.C. and C.W. contributed to scientific discussion. F.T. improved the overview of LncRFM and LncBench. Q.Y. contributed to data statistic and analysis. All authors read the paper, contributed to the discussion and approved the final version.

## Additional information

**Supplementary information.** Supplementary information accompanies this manuscript.

**Correspondence.** Correspondence and requests for materials should be addressed to Heqin Zhu, Peng Xiong or S. Kevin Zhou.

## References

[1] Mattick, J. S. et al. Long non-coding RNAs: definitions, functions, challenges and recommendations. Nature Reviews Molecular Cell Biology 24, 430–447 (2023).

[2] Statello, L., Guo, C.-J., Chen, L.-L. & Huarte, M. Gene regulation by long non-coding RNAs and its biological functions. Nature Reviews Molecular Cell Biology 22, 96–118 (2021).

[3] Lubelsky, Y. & Ulitsky, I. Sequences enriched in Alu repeats drive nuclear localization of long RNAs in human cells. Nature 555, 107–111 (2018).

[4] Yin, Y. et al. U1 snRNP regulates chromatin retention of noncoding RNAs. Nature 580, 147–150 (2020).

[5] Shukla, C. J. et al. High-throughput identification of RNA nuclear enrichment sequences. The EMBO Journal 37, e98452 (2018).

[6] Dominguez, D. et al. Sequence, structure, and context preferences of human RNA binding proteins. Molecular Cell 70, 854–867.e9 (2018).

[7] Somarowthu, S. et al. HOTAIR forms an intricate and modular secondary structure. Molecular Cell 58, 353–361 (2015).

[8] Lu, Z. et al. RNA duplex map in living cells reveals higher-order transcriptome structure. Cell 165, 1267–1279 (2016).

[9] Chen, J. et al. Interpretable RNA foundation model from unannotated data for highly accurate RNA structure and function predictions. arXiv preprint arXiv:2204*.00300* (2022). URL https://arxiv.org/abs/2204.00300.

[10] Penić, R. J., Vlašić, T., Huber, R. G., Wan, Y. & Šikić, M. RiNALMo: general-purpose RNA language models can generalize well on structure prediction tasks. Nature Communications 16, 5671 (2025).

[11] Chen, K. et al. Self-supervised learning on millions of primary RNA sequences from 72 vertebrates improves sequence-based RNA splicing prediction. Briefings in Bioinformatics 25, bbae163 (2024).

[12] Chu, Y. et al. A 5’ UTR language model for decoding untranslated regions of mRNA and function predictions. Nature Machine Intelligence 6, 449–460 (2024).

[13] Romeijn, L., Cats, D., Wolstencroft, K. & Mei, H. LncRNA-BERT: an RNA language model for classifying coding and long non-coding RNA. Connection Science 38, 2624131 (2026).

[14] Wang, N. et al. Multi-purpose RNA language modelling with motif-aware pre-training and type-guided fine-tuning. Nature Machine Intelligence 6, 548–557 (2024).

[15] Zhang, Y. et al. Multiple sequence alignment-based RNA language model and its application to structural inference. Nucleic Acids Research 52, e3 (2024).

[16] Yin, W. et al. ERNIE-RNA: an RNA language model with structure-enhanced representations. Nature Communications 16, 10076 (2025).

[17] Zhu, H. et al. An open structure-guided RNA foundation model for generalizable prediction. Nature (2026). URL 10.1038/s41586-026-11210-2.

[18] Ren, Y. et al. BEACON: benchmark for comprehensive RNA tasks and language models. In Advances in Neural Information Processing Systems (eds Globerson, A. et al.), Vol. 37, 92891–92921 (Curran Associates, Inc., 2024).

[19] Wang, H., Lin, W., Xiao, H., Ye, Q. & Zhang, Y. RNA-Scope: benchmarking RNA language models for RNA sequence understanding. NeurIPS 2025 AI for Science Workshop (2025). URL https://openreview.net/forum?id=lnV3NXpDiC.

[20] Lin, Y., Pan, X. & Shen, H.-B. lncLocator 2.0: a cell-line-specific subcellular localization predictor for long non-coding RNAs with interpretable deep learning. Bioinformatics 37, 2308–2316 (2021).

[21] Zeng, M. et al. DeepLncLoc: a deep learning framework for long non-coding RNA subcellular localization prediction based on subsequence embedding. Briefings in Bioinformatics 23, bbab360 (2022).

[22] Kang, Y.-J. et al. CPC2: a fast and accurate coding potential calculator based on sequence intrinsic features. Nucleic Acids Research 45, W12–W16 (2017).

[23] Han, S. et al. LncFinder: an integrated platform for long non-coding RNA identification utilizing sequence intrinsic composition, structural information and physicochemical property. Briefings in Bioinformatics 20, 2009–2027 (2019).

[24] Tong, X. & Liu, S. CPPred: coding potential prediction based on the global description of RNA sequence. Nucleic Acids Research 47, e43 (2019).

[25] Pan, X., Fan, Y.-X., Yan, J. & Shen, H.-B. IPMiner: hidden ncRNA-protein interaction sequential pattern mining with stacked autoencoder for accurate computational prediction. BMC Genomics 17, 582 (2016).

[26] Xu, Y. et al. PrismNet: predicting protein–RNA interaction using in vivo RNA structural information. Nucleic Acids Research 51, W468–W477 (2023).

[27] Xuan, P., Cao, Y., Zhang, T., Kong, R. & Zhang, Z. Dual convolutional neural networks with attention mechanisms based method for predicting disease-related lncRNA genes. Frontiers in Genetics 10, 416 (2019).

[28] Sheng, N., Cui, H., Zhang, T. & Xuan, P. Attentional multilevel representation encoding based on convolutional and variance autoencoders for lncRNA–disease association prediction. Briefings in Bioinformatics 22, bbaa067 (2021).

[29] Zhu, H. et al. Deep generalizable prediction of RNA secondary structure via base pair motif energy. Nature Communications 16, 5856 (2025).

30. Zhu, H., et al. NC-Bench and NCfold: a benchmark and closed-loop framework for RNA non-canonical base-pair prediction. In The Fourteenth International Conference on Learning Representations (2026).

31. Valhalla Team. Leveraging secondary-structure information for accurate nucleic acid structure prediction with OFoldNA. arXiv (2026). URL https://arxiv.org/abs/2609.37414.

32. Chang, A., et al. ReTIF: granularity-aware multitask interaction routing for RNA–compound interaction prediction and binding-site localization. bioRxiv (2026). URL https://www.biorxiv.org/content/early/2026/09/06/2026.09.02.748859.

[33] Zhang, F. et al. IRESeek: structure-informed deep learning method for accurate identification of internal ribosome entry sites in circular RNAs. NAR Genomics and Bioinformatics 7, lqaf210 (2025).

34. Gu, A. & Dao, T. Mamba: linear-time sequence modeling with selective state spaces. In First Conference on Language Modeling (2024). URL https://openreview.net/forum?id=tEYskw1VY2.

[35] Zhao, W. et al. POSTAR3: an updated platform for exploring post-transcriptional regulation coordinated by RNA-binding proteins. Nucleic Acids Research 50, D287–D294 (2022).

[36] Mas-Ponte, D. et al. LncATLAS database for subcellular localization of long noncoding RNAs. RNA 23, 1080–1087 (2017).

[37] Li, Z. et al. LncBook 2.0: integrating human long non-coding RNAs with multiomics annotations. Nucleic Acids Research 51, D186–D191 (2023).

[38] Zubradt, M. et al. DMS-MaPseq for genome-wide or targeted RNA structure probing in vivo. Nature Methods 14, 75–82 (2017).

[39] Huang, L. et al. LinearFold: linear-time approximate RNA folding by 5’-to-3’ dynamic programming and beam search. Bioinformatics 35, i295–i304 (2019).

40. Zhu, H., et al. NACraft: programmatic nucleic-acid aptamer design via all-atom structure-model feedback. bioRxiv (2026). URL https://www.biorxiv.org/content/early/2026/08/18/2026.08.15.744087.

41. Valhalla Team. De novo design of monoclonal and bispecific antibodies with OFAntibody. arXiv (2026). URL https://arxiv.org/abs/2610.09548.

[42] RNAcentral Consortium. RNAcentral 2021: secondary structure integration, improved sequence search and new member databases. Nucleic Acids Research 49, D212–D220 (2021).

[43] Su, Z.-D. et al. iLoc-lncRNA: predict the subcellular location of lncRNAs by incorporating octamer composition into general PseKNC. Bioinformatics 34, 4196–4204 (2018).

[44] Bai, T. & Liu, B. ncRNALocate-EL: a multi-label ncRNA subcellular locality prediction model based on ensemble learning. Briefings in Functional Genomics 22, 442–452 (2023).

[45] Frankish, A. et al. GENCODE reference annotation for the human and mouse genomes. Nucleic Acids Research 47, D766–D773 (2019).

[46] O’Leary, N. A. et al. Reference sequence (RefSeq) database at NCBI: current status, taxonomic expansion, and functional annotation. Nucleic Acids Research 44, D733–D745 (2016).

[47] Tate, J. G. et al. COSMIC: the catalogue of somatic mutations in cancer. Nucleic Acids Research 47, D941–D947 (2019).

[48] Landrum, M. J. et al. ClinVar: improving access to variant interpretations and supporting evidence. Nucleic Acids Research 46, D1062–D1067 (2018).

[49] Li, G. et al. HydraRNA: a hybrid architecture based full-length RNA language model. Genome Biology 26, 383 (2025).

[50] Li, W. & Godzik, A. CD-HIT: a fast program for clustering and comparing large sets of protein or nucleotide sequences. Bioinformatics 22, 1658–1659 (2006).

[51] Devlin, J., Chang, M.-W., Lee, K. & Toutanova, K. BERT: pre-training of deep bidirectional transformers for language understanding. In Proceedings of the 2019 Conference of the North American Chapter of the Association for Computational Linguistics: Human Language Technologies, Volume 1 (Long and Short Papers) (eds Burstein, J., Doran, C. & Solorio, T.), 4171–4186 (Association for Computational Linguistics, 2019).

[52] Houlsby, N. et al. Parameter-efficient transfer learning for NLP. In Proceedings of the 36th International Conference on Machine Learning (eds Chaudhuri, K. & Salakhutdinov, R.), Vol. 97 of Proceedings of Machine Learning Research, 2790–2799 (PMLR, 2019).

[53] Paszke, A. et al. PyTorch: an imperative style, high-performance deep learning library. In Advances in Neural Information Processing Systems (eds Wallach, H. M. et al.), Vol. 32, 8026–8037 (Curran Associates, Inc., 2019).

54. Loshchilov, I. & Hutter, F. Decoupled weight decay regularization. Published as a conference paper at the International Conference on Learning Representations (2019). URL https://openreview.net/forum?id=Bkg6RiCqY7.

[55] Gneiting, T. & Raftery, A. E. Strictly proper scoring rules, prediction, and estimation. Journal of the American Statistical Association 102, 359–378 (2007).

[56] Efron, B. Bootstrap methods: another look at the jackknife. The Annals of Statistics 7, 1–26 (1979).

[57] Chang, A. et al. LncCorpus. Zenodo [Data set] (2026). URL 10.5281/zenodo.24344559.

[58] Chang, A. et al. LncBench. Zenodo [Data set] (2026). URL 10.5281/zenodo.24345986.

