## Supplemental file for "An open long non-coding RNA foundation model for generalizable prediction"

#### 1 Extended Data Figures and Tables

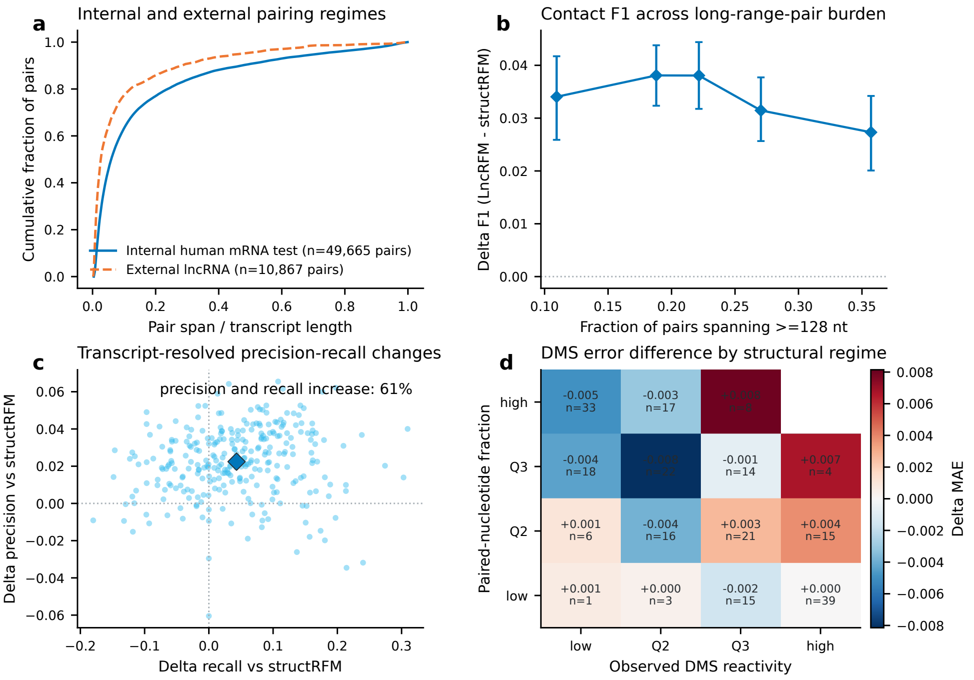


**Extended Data Fig. 1**: **Contact and DMS prediction across pairing contexts. a**, Cumulative relative pair-span distributions for internal mRNA and external lncRNA ($n = 49,665$ and 10,867 pairs). **b**, Mean contact ΔF1 across quintiles of the fraction of pairs spanning $\geq 128$ nt ($n = 56$–60 transcripts/bin; 95% intervals, 2,000 transcript bootstraps). **c**, Per-transcript precision/recall differences ($n = 291$). **d**, Mean DMS ΔMAE by pairing density and reactivity ($n = 232$; bin counts shown). Differences are LncRFM minus structRFM; negative ΔMAE favours LncRFM.

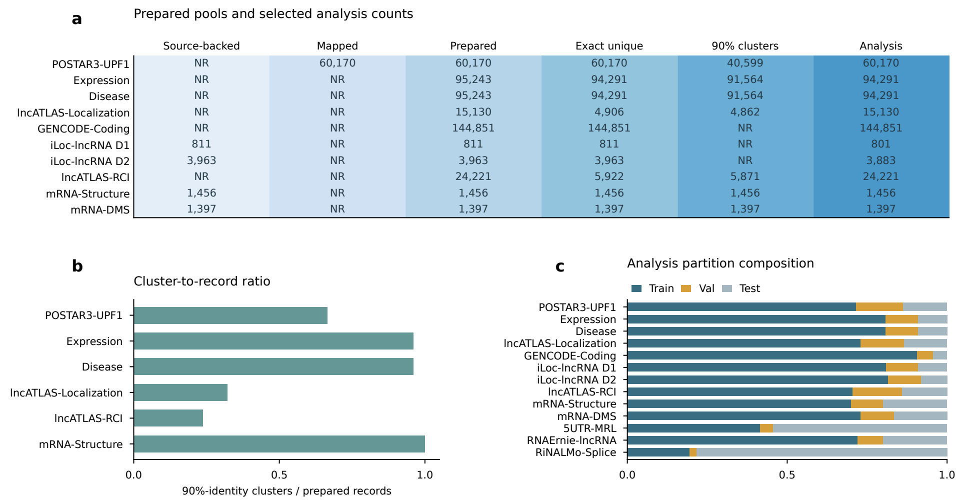


**Extended Data Fig. 2**: **Dataset preparation stages and analysis-partition composition. a**, Documented source-backed, identifier-mapped, prepared-record, exact-unique-sequence, 90%-cluster and analysis counts across datasets. NR denotes unverified upstream counts. **b**, Ratio of 90%-identity clusters to prepared records: Expression and Disease use MMseqs Linclust followed by exact-sequence merging; other documented cluster counts use CD-HIT. **c**, Train/Val/Test proportions for analysis (partition counts in Extended Data Table 1).

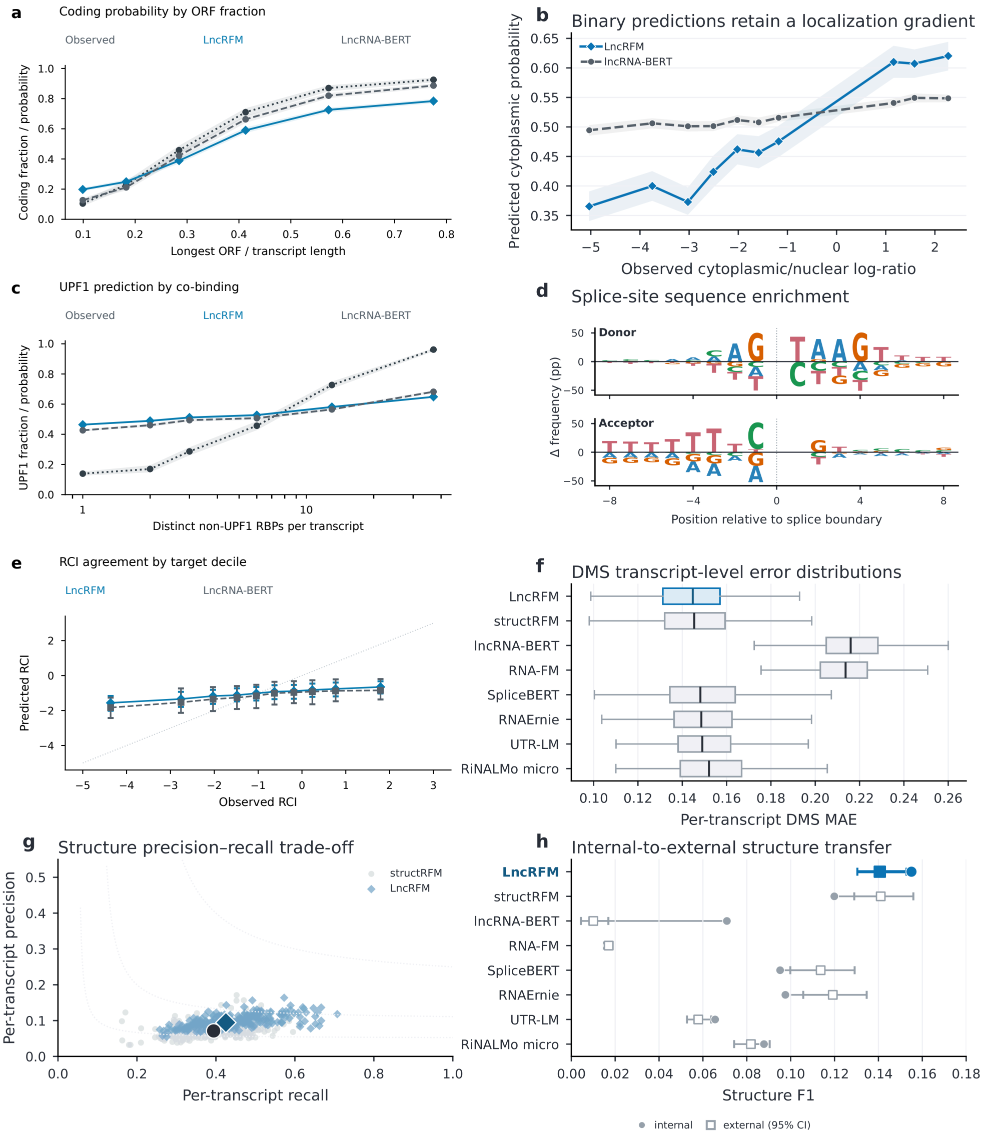


**Extended Data Fig. 3**: **Score gradients and transcript-level error distributions. a–c**, Predicted probabilities across longest-ORF fraction, cytoplasmic/nuclear log-ratio and non-UPF1 RBP diversity (6,316, 2,031 and 8,216 records, respectively); observed positive fractions are also shown in **a,c**. Bands, 95% intervals from 2,000 within-bin record bootstraps for predictions and Wilson intervals for observed fractions; shading in **a,c**, observation–prediction gaps. **d**, Positive-minus-negative splice-flank nucleotide frequencies. **e**, Mean predicted versus observed RCI across deciles (3,411 records); bars show prediction interquartile ranges (IQRs), and the diagonal marks equality. **f**, DMS MAE for 232 transcripts (boxes, IQRs; lines, medians; whiskers, extremes within 1.5 × IQR of box limits; outliers omitted). **g**, Contact precision/recall for 291 transcripts (outlined symbols, medians; dotted curves, iso-F1). **h**, Internal/external contact F1; external 95% intervals use 2,000 transcript bootstraps ($n = 30$). LncRFM is blue.

**Extended Data Table 1**: Overview and statistics of the 15 LncBench datasets across 10 prediction tasks. Train, Val and Test give the evaluated partition record counts across all adaptation and external generalization benchmarks.

| Category | Dataset | Prediction task | Task type | Source | Train | Val | Test | Mean length (nt) |
|---|---|---|---|---|---|---|---|---|
| **Internal adaptation: 6 core lncRNA tasks (9 datasets)** | | | | | | | | |
| Localization | iLoc-lncRNA D1 | Subcellular localization | Multilabel | iLoc-lncRNA [59] | 648 | 81 | 72 | 2,470.4 |
| Localization | iLoc-lncRNA D2 | Subcellular localization | Multilabel | NCRNALocate-EL [60] | 3,170 | 396 | 317 | 3,819.2 |
| Localization | lncATLAS-Localization | Subcellular localization (nuclear–cytoplasmic) | Binary | lncATLAS [61] | 11,034 | 2,065 | 2,031 | 2,766.9 |
| Localization | lncATLAS-RCI | Cytoplasmic-to-nuclear RCI | Regression | lncATLAS [61] | 17,083 | 3,727 | 3,411 | 2,729.2 |
| Coding | GENCODE-Coding | Coding potential | Binary | GENCODE [62], RefSeq [63] | 131,264 | 7,271 | 6,316 | 2,459.2 |
| Coding | RNAErnie-lncRNA | Coding potential | Binary | RNAErnie [64] | 115,542 | 12,838 | 32,096 | 1,343.5 |
| Variation | Disease | Official variation-record status | Binary | LncBook 2.0 [65] | 76,196 | 9,524 | 8,571 | 1,975.4 |
| Context | Expression | Expression capacity | Multilabel | LncBook 2.0 [65] | 76,196 | 9,524 | 8,571 | 1,969.7 |
| Interaction | POSTAR3-UPF1 | RNA–protein binding | Binary | POSTAR3 [66], GENCODE [62] | 43,078 | 8,876 | 8,216 | 1,239.6 |
| **Internal adaptation: 4 structural and regulatory assay tasks (4 datasets)** | | | | | | | | |
| Structure | mRNA-Structure | RNA contact | Pairwise binary | Rouskinlab [67] | 1,019 | 146 | 291 | 647.3 |
| Structure | mRNA-DMS | DMS reactivity | Per-base regression | Rouskinlab [67] | 1,019 | 146 | 232 | 647.3 |
| Translation | 5UTR-MRL | Mean ribosome load (MRL) | Regression | Optimus 5-Prime [68] | 76,319 | 7,600 | 100,017 | 55.2 |
| Splicing | RiNALMo-Splice | Splice-site recognition (donor) | Binary | RiNALMo [69] | 19,775 | 2,198 | 80,000 | 400.0 |
| | | Splice-site recognition (acceptor) | Binary | RiNALMo [69] | 19,961 | 2,218 | 80,000 | 400.0 |
| **External OOD generalization: 2 transfer tasks (2 datasets; evaluation only)** | | | | | | | | |
| Coding | RNAChallenge | Coding potential | Binary | RNAChallenge | – | – | 27,283 | 1,635.2 |
| Structure | lncRNA-Structure | RNA contact (lncRNA) | Pairwise binary | Rouskinlab [67] | – | – | 30 | 1,298.3 |
| **Total (10 tasks; 15 datasets)** | | | | | **592,304** | **66,610** | **357,454** | – |

Mean lengths (nt) summarize the evaluated partition records across tasks. RiNALMo-Splice comprises donor and acceptor splice-site recognition subtasks; both enter totals, which count records separately for each subtask.

| Model | Parameters (M) | Tokenizer | Context (tokens) | Trainable (M) | Trainable (%) |
|---|---|---|---|---|---|
| LncRFM | 48.960 | nucleotide | 2050 | 0.825 | 1.7 |
| structRFM | 86.876 | released nucleotide | 512 | 0.825 | 0.9 |
| LncRNA-BERT | 85.934 | non-overlapping 3-mer | 768 | 0.825 | 1.0 |
| RNA-FM | 99.853 | released nucleotide | 1024 | 0.743 | 0.7 |
| SpliceBERT | 20.379 | released nucleotide | 510 | 0.660 | 3.2 |
| RNAErnie | 86.889 | released nucleotide | 512 | 0.825 | 0.9 |
| UTR-LM | 1.622 | released nucleotide | 1024 | 0.412 | 25.4 |
| RiNALMo micro | 34.122 | released nucleotide | 1024 | 0.639 | 1.9 |

**Extended Data Table 2**: Model sizes, context limits and trainable parameters for the binary adapter/head reference configuration. Counts include the encoder, adapter and instantiated binary head; they are not asserted to be identical for multilabel and regression heads. Context is measured in encoded tokens; LncRNA-BERT uses non-overlapping 3-mers and the other seven models use nucleotide tokens.

| Dataset | Prediction head | Loss | Learning rate | Early-stopping metric |
|---|---|---|---|---|
| POSTAR3-UPF1 | sequence adapter + binary MLP | weighted BCE | $1 \times 10^{-5}$ | validation F1 |
| Expression | sequence adapter + five-logit MLP | BCE | $5 \times 10^{-5}$ | validation micro-F1 |
| Disease | sequence adapter + binary MLP | weighted BCE | $1 \times 10^{-4}$ | validation F1 |
| lncATLAS-Localization | sequence adapter + binary MLP | weighted BCE | $5 \times 10^{-5}$ | validation F1 |
| GENCODE-Coding | sequence adapter + binary MLP | weighted BCE | $1 \times 10^{-5}$ | validation F1 |
| iLoc-lncRNA D1 | sequence adapter + multilabel MLP | BCE | $1 \times 10^{-4}$ | validation micro-F1 |
| iLoc-lncRNA D2 | sequence adapter + multilabel MLP | BCE | $3 \times 10^{-5}$ | validation micro-F1 |
| lncATLAS-RCI | sequence adapter + scalar MLP | MSE | $1 \times 10^{-5}$ | validation Pearson $r$ |
| mRNA-Structure | token adapter + symmetric Q–K contact head | weighted BCE | $1 \times 10^{-4}$ | validation F1 |
| mRNA-DMS | token adapter + nucleotide regressor | MSE | $1 \times 10^{-4}$ | validation Pearson $r$ |
| 5UTR-MRL | sequence adapter + scalar MLP | MSE | $5 \times 10^{-5}$ | Validation squared Pearson correlation ($r^2$) |
| RNAErnie-lncRNA | sequence adapter + binary MLP | weighted BCE | $5 \times 10^{-5}$ | validation ROC-AUC |
| RiNALMo-Splice donor | sequence adapter + binary MLP | weighted BCE | $5 \times 10^{-5}$ | validation F1 |
| RiNALMo-Splice acceptor | sequence adapter + binary MLP | weighted BCE | $5 \times 10^{-5}$ | validation F1 |

**Extended Data Table 3**: Task heads, losses, learning rates and validation checkpoint-selection metrics across LncBench adaptation tasks. For POSTAR3-UPF1, Expression, Disease, lncATLAS-RCI and GENCODE-Coding, LncRFM used effective batch size 32 (microbatch 16 with two-step accumulation, or microbatch 4 with eight-step accumulation for GENCODE-Coding), early-stopping patience 5 and minimum improvement 0.001, with training capped at 100 epochs (40 epochs for GENCODE-Coding). Other adaptation tasks used a 15-epoch minimum and patience 10, except the splice classifiers (five fixed epochs). All models were evaluated using the checkpoint achieving optimal validation performance under their respective task configurations.

| Dataset | Source distribution / version | Species | Unit | Sequence source |
|---|---|---|---|---|
| LncCorpus | RNAcentral hybrid corpus; MUSES/LinearFold pairing annotations | Not recorded | RNA sequence | RNAcentral |
| iLoc-lncRNA D1 | iLoc-lncRNA precompiled table | Human | RNA record | iLoc-lncRNA table |
| iLoc-lncRNA D2 | NCRNALocate-EL benchmark and independent FASTA collections | Human | RNA record | Benchmark and independent FASTA |
| lncATLAS-Localization | lncATLAS; GENCODE v47 | Human | Gene–cell line record | GENCODE v47 |
| lncATLAS-RCI | lncATLAS; GENCODE v47 | Human | Gene–cell line record | GENCODE v47 |
| GENCODE-Coding | GENCODE/RefSeq; expanded processed splits | Human | Transcript | GENCODE/RefSeq |
| RNAErnie-lncRNA | Processed downstream collection (Methods Section 4.1) | Human/mouse | Transcript | RNAErnie data |
| Disease | LncBook 2.0; official variation | Human | Gene | LncBook 2.0 official GTF |
| Expression | LncBook 2.0; official expression | Human | Gene | LncBook 2.0 official GTF |
| POSTAR3-UPF1 | POSTAR3; GENCODE v47 | Human | Transcript | GENCODE v47 |
| mRNA-Structure | Processed Rouskinlab collection | Human | Transcript | Rouskinlab |
| mRNA-DMS | Processed Rouskinlab collection | Human | Transcript | Rouskinlab |
| 5UTR-MRL | Processed downstream collection (Methods Section 4.1) | Mixed | 5' UTR | Optimus 5-Prime |
| RiNALMo-Splice | Processed downstream collection (Methods Section 4.1) | Multi-species | 400-nt window | RiNALMo data |
| RNAChallenge | RNAChallenge FASTA collection | Multi-species (animal, plant, fungi) | Transcript | RNAChallenge |
| lncRNA-Structure | Processed Rouskinlab collection | Human | Transcript | Rouskinlab |

**Extended Data Table 4**: Upstream data sources, versions and statistical units. Entries identify upstream databases or processed benchmark collections. Expression and Disease use official LncBook gene records represented by the longest transcript in the official GTF. Expression encodes five expression-capacity contexts (HC/MC/LC versus NE). Disease records the presence of any official variation annotation, including dbSNP, COSMIC, ClinVar or GWAS records.

| Dataset | Raw/source-backed | Mapped | Benchmark records | Exact-unique | 90% clusters |
|---|---|---|---|---|---|
| LncCorpus | 31,121,367 | NA | 24,771,981 | NA | NA |
| iLoc-lncRNA D1 | 801 | NA | 801 | 801 | NA |
| iLoc-lncRNA D2 | 3,883 | NA | 3,883 | 3,883 | NA |
| lncATLAS-Localization | — | — | 15,130 | 4,906 | 4,862 |
| lncATLAS-RCI | NR | NR | 24,221 | 5,922 | 5,871 |
| GENCODE-Coding | NR | NR | 144,851 | 144,851 | NR |
| RNAErnie-lncRNA | — | NA | 160,476 | 159,748 | NA |
| Disease | NR | — | 94,291 | 94,291 | 91,564 |
| Expression | NR | — | 94,291 | 94,291 | 91,564 |
| POSTAR3-UPF1 | NR | 60,170 | 60,170 | 60,170 | 40,599 |
| mRNA-Structure | 1,456 | NA | 1,456 | 1,456 | 1,456 |
| mRNA-DMS | 1,397 | NA | 1,397 | 1,397 | 1,397 |
| 5UTR-MRL | — | NA | 183,936 | 183,936 | NA |
| RiNALMo-Splice | — | NA | 204,152 | 204,057 | NA |
| RNAChallenge | 27,283 | NA | 27,283 | 27,283 | NA |
| lncRNA-Structure | 30 | NA | 30 | 24 | NA |

**Extended Data Table 5**: Record counts across dataset preparation stages. Raw/source-backed is the earliest documented count; Benchmark records denotes the finalized records across the Train, Validation and Test partitions (matching Extended Data Table 1; for LncCorpus, 24,648,121 training and 123,860 validation records). Exact-unique and 90% clusters summarize unique sequence and 90%-identity cluster counts (using CD-HIT where recorded, except Expression and Disease, which use MMseqs Linclust followed by exact-sequence merging). NA, not applicable; NR or —, not recorded.

**Extended Data Table 6:** Primary-metric performance across 15 LncBench datasets and 10 prediction tasks.

| Task | Dataset | N | Metric | Method | Score |
|---|---|---|---|---|---|
| RNA–protein binding prediction | POSTAR3-UPF1 | 8,216 | AUROC ↑ | structRFM | 0.714 |
| | | | | LncRNA-BERT | 0.714 |
| | | | | RNA-FM | 0.630 |
| | | | | SpliceBERT | 0.626 |
| | | | | RNAErnie | 0.623 |
| | | | | UTR-LM | 0.609 |
| | | | | RiNALMo micro | 0.608 |
| | | | | LncRFM | **0.757** |
| Expression capacity prediction | Expression | 8,571 | Macro-AUPRC ↑ | structRFM | 0.465 |
| | | | | LncRNA-BERT | 0.470 |
| | | | | RNA-FM | 0.432 |
| | | | | SpliceBERT | 0.423 |
| | | | | RNAErnie | 0.421 |
| | | | | UTR-LM | 0.422 |
| | | | | RiNALMo micro | 0.424 |
| | | | | LncRFM | **0.498** |
| Variation-record status prediction | Disease | 8,571 | AUROC ↑ | structRFM | 0.661 |
| | | | | LncRNA-BERT | 0.740 |
| | | | | RNA-FM | 0.646 |
| | | | | SpliceBERT | 0.627 |
| | | | | RNAErnie | 0.630 |
| | | | | UTR-LM | 0.593 |
| | | | | RiNALMo micro | 0.651 |
| | | | | LncRFM | **0.791** |
| Subcellular localization | lncATLAS-Localization | 2,031 | F1 ↑ | structRFM | 0.472 |
| | | | | LncRNA-BERT | 0.520 |
| | | | | RNA-FM | 0.481 |
| | | | | SpliceBERT | 0.460 |
| | | | | RNAErnie | 0.489 |
| | | | | UTR-LM | 0.444 |
| | | | | RiNALMo micro | 0.455 |
| | | | | LncRFM | **0.583** |
| | iLoc-lncRNA D1 | 72 | Micro-F1 ↑ | structRFM | 0.729 |
| | | | | LncRNA-BERT | 0.743 |
| | | | | RNA-FM | 0.727 |
| | | | | SpliceBERT | 0.721 |
| | | | | RNAErnie | 0.712 |
| | | | | UTR-LM | 0.702 |
| | | | | RiNALMo micro | 0.727 |
| | | | | LncRFM | **0.764** |
| | iLoc-lncRNA D2 | 317 | Micro-F1 ↑ | structRFM | 0.743 |
| | | | | LncRNA-BERT | 0.767 |
| | | | | RNA-FM | 0.756 |
| | | | | SpliceBERT | 0.757 |
| | | | | RNAErnie | 0.756 |
| | | | | UTR-LM | 0.752 |
| | | | | RiNALMo micro | 0.762 |
| | | | | LncRFM | **0.770** |

**Extended Data Table 6 (continued)**

| Task | Dataset | N | Metric | Method | Score |
|---|---|---|---|---|---|
| Coding potential prediction | GENCODE-Coding | 6,316 | AUROC ↑ | structRFM | 0.911 |
| | | | | LncRNA-BERT | 0.968 |
| | | | | RNA-FM | 0.818 |
| | | | | SpliceBERT | 0.806 |
| | | | | RNAErnie | 0.806 |
| | | | | UTR-LM | 0.796 |
| | | | | RiNALMo micro | 0.821 |
| | | | | LncRFM | **0.980** |
| | RNAErnie-lncRNA | 32,096 | AUROC ↑ | structRFM | 0.919 |
| | | | | LncRNA-BERT | **0.942** |
| | | | | RNA-FM | 0.744 |
| | | | | SpliceBERT | 0.727 |
| | | | | RNAErnie | 0.722 |
| | | | | UTR-LM | 0.741 |
| | | | | RiNALMo micro | 0.750 |
| | | | | LncRFM | 0.932 |
| | RNAChallenge | 27,283 | F1 ↑ | structRFM | 0.396 |
| | | | | LncRNA-BERT | 0.237 |
| | | | | RNA-FM | **0.429** |
| | | | | SpliceBERT | 0.315 |
| | | | | RNAErnie | 0.399 |
| | | | | UTR-LM | 0.353 |
| | | | | RiNALMo micro | 0.273 |
| | | | | LncRFM | 0.374 |
| Relative concentration index prediction | lncATLAS-RCI | 3,411 | Pearson r ↑ | structRFM | 0.311 |
| | | | | LncRNA-BERT | 0.344 |
| | | | | RNA-FM | 0.175 |
| | | | | SpliceBERT | 0.179 |
| | | | | RNAErnie | 0.180 |
| | | | | UTR-LM | 0.193 |
| | | | | RiNALMo micro | 0.180 |
| | | | | LncRFM | **0.403** |
| RNA contact prediction | mRNA-Structure | 291 | F1 ↑ | structRFM | 0.120 |
| | | | | LncRNA-BERT | 0.071 |
| | | | | RNA-FM | 0.017 |
| | | | | SpliceBERT | 0.095 |
| | | | | RNAErnie | 0.098 |
| | | | | UTR-LM | 0.066 |
| | | | | RiNALMo micro | 0.088 |
| | | | | LncRFM | **0.155** |
| | lncRNA-Structure | 30 | F1 ↑ | structRFM | **0.141** |
| | | | | LncRNA-BERT | 0.010 |
| | | | | RNA-FM | 0.017 |
| | | | | SpliceBERT | 0.114 |
| | | | | RNAErnie | 0.119 |
| | | | | UTR-LM | 0.058 |
| | | | | RiNALMo micro | 0.082 |
| | | | | LncRFM | 0.141 |

**Extended Data Table 6 (continued)**

| Task | Dataset | N | Metric | Method | Score |
|---|---|---|---|---|---|
| DMS reactivity prediction | mRNA-DMS | 232 | Pearson r ↑ | structRFM | 0.659 |
| | | | | LncRNA-BERT | 0.075 |
| | | | | RNA-FM | 0.087 |
| | | | | SpliceBERT | 0.639 |
| | | | | RNAErnie | 0.638 |
| | | | | UTR-LM | 0.647 |
| | | | | RiNALMo micro | 0.655 |
| | | | | LncRFM | **0.669** |
| Mean ribosome load prediction | 5UTR-MRL | 100,017 | Pearson $r^2$ ↑ | structRFM | 0.273 |
| | | | | LncRNA-BERT | 0.277 |
| | | | | RNA-FM | 0.146 |
| | | | | SpliceBERT | 0.081 |
| | | | | RNAErnie | 0.124 |
| | | | | UTR-LM | 0.121 |
| | | | | RiNALMo micro | 0.149 |
| | | | | LncRFM | **0.354** |
| Splice-site recognition | RiNALMo-Splice | 80,000 | Mean F1 ↑ | structRFM | 0.687 |
| | | | | LncRNA-BERT | 0.660 |
| | | | | RNA-FM | **0.707** |
| | | | | SpliceBERT | 0.667 |
| | | | | RNAErnie | 0.668 |
| | | | | UTR-LM | 0.636 |
| | | | | RiNALMo micro | 0.635 |
| | | | | LncRFM | 0.641 |

### 2 Supplementary Methods

#### 2.1 Coding Potential Prediction

`GENCODE-Coding` distinguishes human lncRNA from protein-coding transcripts using a curated GENCODE [62] and RefSeq [63] transcript resource comprising $131,264$ training, $7,271$ validation and $6,316$ test transcript records ($144,851$ total constructed records; Extended Data Tables 1 and 5). Separately, the external RNAChallenge benchmark was evaluated without further tuning using the checkpoint adapted on the human GENCODE-Coding splits.

#### 2.2 Subcellular Localization Prediction

***iLoc-lncRNA D1.***

iLoc-lncRNA D1 was obtained from a precompiled multilabel lncRNA localization table (811 initial records) [59] with aligned identifiers, sequences and compartment labels. After standardizing field names, retaining lncRNA records and excluding 10 records lacking valid subcellular localization labels (801 retained), we encoded each record's supplied compartment annotations as a multihot target.

***iLoc-lncRNA D2.***

iLoc-lncRNA D2 uses human lncRNA records from the NCRNALocate-EL benchmark and independent FASTA collections [60]. We parsed identifiers and compartment annotations from FASTA headers, combined the collections, removed exact identifier–sequence duplicates and disambiguated repeated identifiers. Standardized compartment sets serve as multilabel targets for the retained sequences.

#### 2.3 lncATLAS Localization Tasks

The lncATLAS localization table gives gene–cell-line fractionation measurements [61], while the model requires transcript sequences. We selected non-coding entries, normalized Ensembl IDs and attached the longest GENCODE v47 lncRNA transcript per gene [62], retaining individual gene–cell-line records ($24,221$ constructed records across $5,922$ exact-unique sequences and $5,871$ clusters at 90% sequence identity for lncATLAS-RCI). For binary lncATLAS-Localization, `ratio2` is $\log_2$(cytosol/nucleus); values $\geq 1$ indicate cytoplasmic enrichment (label 1) and values $\leq -1$ nuclear enrichment (label 0). Intermediate records were excluded. All records from a given gene remained in one partition.

lncATLAS-RCI uses the same sequence attachment and retains the continuous cytoplasmic-to-nuclear RCI value ($17,083$ training, $3,727$ validation and $3,411$ test records).

#### 2.4 LncBook disease and expression-context annotations

`Disease` and `Expression` were constructed from official LncBook 2.0 human lncRNA gene annotations [65], with each gene represented by its longest transcript sequence in the official GRCh38 GTF. Both tasks share a prepared pool of $95,243$ official gene records (94,291 exact-unique sequences), clustered at 90% sequence identity using MMseqs Linclust followed by merging of clusters sharing identical or reverse-complement sequences (91,564 final clusters; Extended Data Table 5). Each task was partitioned into 76,196 training, 9,524 validation and 8,571 test gene records (the two tasks do not share identical test gene lists, whereas all eight models were evaluated on the exact same test IDs within each task).

For `Disease`, a binary target was defined as 1 if a gene had at least one recorded entry in the official LncBook variation table (incorporating records from dbSNP, COSMIC [70], ClinVar [71] and GWAS) and 0 otherwise; positive labels therefore denote documented official variation annotations rather than verified disease causality. For `Expression`, five official expression-capacity contexts (*normal_tissue_cell_line*, *organ_development*, *preimplantation_embryo*, *cell_differentiation* and *subcellular_localization*) were binarized by encoding high, medium or low expression capacity (HC, MC or LC) as 1 and no expression (NE) as 0, yielding a five-dimensional multilabel target. Records with multiple positive contexts retained all corresponding positive labels, and records with NE across all five contexts retained an all-zero target vector.

#### 2.5 lncRNA–RBP Interaction Prediction

We converted human POSTAR3 binding peaks [66] to transcript-level targets using GENCODE v47 lncRNA exon coordinates and sequences [62] (61,083 constructed transcript records across 60,170 exact-unique sequences and 40,599 CD-HIT clusters at 90% identity; 43,078 training, 8,876 validation and 8,216 test transcripts). Exons were indexed from the lncRNA GTF, and peaks were matched by chromosome, strand and interval overlap. Generic `RBP_occupancy` entries were discarded. For each transcript, the resulting target lists RBPs with mapped peak support, with UPF1 defining the binary classification label.

Co-binding diversity was calculated as the number of distinct non-UPF1 RBPs annotated for each transcript in the complete 216-RBP annotation matrix.

#### 2.6 Supplementary Structure-Oriented Tasks

The processed Rouskinlab `human_mRNA` resource pairs sequences with secondary-structure annotations and DMS-MaPseq measurements [67]. We uppercased sequences, converted T to U and exported two views: mRNA-Structure for contact prediction and mRNA-DMS for nucleotide reactivity.

CD-HIT-EST clusters defined a shared, approximately 7:1:2 source training/validation/test partition for both views [72]. The 30-record Rouskinlab lncRNA collection formed the external lncRNA-Structure set and was evaluated with the internally selected mRNA-Structure checkpoint.

#### 2.7 Transfer Tasks

Sequences in the three transfer datasets (Methods Section 4.1) were uppercased and checked for compatibility with the downstream tokenizer's RNA/DNA alphabet.

##### *5' UTR mean ribosome load regression.*

`5UTR-MRL` predicts mean ribosome load from 5' UTR sequences using measurements from the massively parallel translation assay used to develop Optimus 5-Prime [68]. Following the processed benchmark collection of ref. [73], we used 76,319 training and 7,600 validation records, and evaluated on the full 100,017-record main test split (combining the 7,600 human endogenous 5' UTR test sequences highlighted in ref. [73] with 92,417 synthetic random 5' UTR test sequences from the Optimus 5-Prime assay [68]). Sequences span 25–100 nt, with a mean of 55.2 nt across these partitions (Extended Data Table 1).

##### *lncRNA-versus-protein-coding transfer classification.*

RNAErnie-lncRNA distinguishes lncRNAs from protein-coding RNAs. Its 115,542 training, 12,838 validation and 32,096 test records comprise 80,238 sequences per class overall. Lengths range from 8 to 6,144 nt, with a mean of 1,343.5 nt.

##### *Splice donor and acceptor recognition.*

RiNALMo-Splice contains separate donor and acceptor binary tasks on 400-nt windows. Donor recognition has 19,775/2,198/80,000 training/validation/test records, and acceptor recognition has 19,961/2,218/80,000. Both are approximately class-balanced. The 13-dataset comparison treats the two tasks as one endpoint using their mean F1.

### Supplementary References


[59] Su, Z.-D. *et al.* iLoc-lncRNA: predict the subcellular location of lncRNAs by incorporating octamer composition into general PseKNC. *Bioinformatics* **34**, 4196–4204 (2018).

[60] Bai, T. & Liu, B. ncRNALocate-EL: a multi-label ncRNA subcellular locality prediction model based on ensemble learning. *Briefings in Functional Genomics* **22**, 442–452 (2023).

[61] Mas-Ponte, D. *et al.* LncATLAS database for subcellular localization of long noncoding RNAs. *RNA* **23**, 1080–1087 (2017).

[62] Frankish, A. *et al.* GENCODE reference annotation for the human and mouse genomes. *Nucleic Acids Research* **47**, D766–D773 (2019).

[63] O'Leary, N. A. *et al.* Reference sequence (RefSeq) database at NCBI: current status, taxonomic expansion, and functional annotation. *Nucleic Acids Research* **44**, D733–D745 (2016).

[64] Wang, N. *et al.* Multi-purpose RNA language modelling with motif-aware pre-training and type-guided fine-tuning. *Nature Machine Intelligence* **6**, 548–557 (2024).

[65] Li, Z. *et al.* LncBook 2.0: integrating human long non-coding RNAs with multi-omics annotations. *Nucleic Acids Research* **51**, D186–D191 (2023).

[66] Zhao, W. *et al.* POSTAR3: an updated platform for exploring post-transcriptional regulation coordinated by RNA-binding proteins. *Nucleic Acids Research* **50**, D287–D294 (2022).

[67] Zubradt, M. *et al.* DMS-MaPseq for genome-wide or targeted RNA structure probing in vivo. *Nature Methods* **14**, 75–82 (2017).

[68] Sample, P. J. *et al.* Human 5' UTR design and variant effect prediction from a massively parallel translation assay. *Nature Biotechnology* **37**, 803–809 (2019).

[69] Penić, R. J., Vlašić, T., Huber, R. G., Wan, Y. & Šikić, M. RiNALMo: general-purpose RNA language models can generalize well on structure prediction tasks. *Nature Communications* **16**, 5671 (2025).

[70] Tate, J. G. *et al.* COSMIC: the catalogue of somatic mutations in cancer. *Nucleic Acids Research* **47**, D941–D947 (2019).

[71] Landrum, M. J. *et al.* ClinVar: improving access to variant interpretations and supporting evidence. *Nucleic Acids Research* **46**, D1062–D1067 (2018).

[72] Li, W. & Godzik, A. CD-HIT: a fast program for clustering and comparing large sets of protein or nucleotide sequences. *Bioinformatics* **22**, 1658–1659 (2006).

[73] Li, G. *et al.* HydraRNA: a hybrid architecture based full-length RNA language model. *Genome Biology* **26**, 383 (2025).
